# The critical role of isomiRs in accurate differential expression analysis of miRNA-seq data

**DOI:** 10.1101/2024.03.28.587190

**Authors:** Eloi Schmauch, Yassine Attia, Pia Laitinen, Tiia A. Turunen, Piia Bartos, Mari-Anna Vaananen, Tarja Malm, Pasi Tavi, Manolis Kellis, Minna U Kaikkonen, Suvi Linna-Kuosmanen

## Abstract

MicroRNAs (miRNAs) are crucial for the regulation of gene expression and are promising biomarkers and therapeutic targets. miRNA isoforms (isomiRs) differ in their start/end offsets, which can impact the target gene selection and non-canonical function of the miRNA species. In addition, isomiRs frequently differ in their expression patterns from their parent miRNAs, yet their roles and tissue-specific responses are currently understudied, leading to their typical omission in miRNA research. Here, we evaluate the expression differences of isomiRs across conditions and their impact on standard miRNA-seq quantification results. We analyze 28 public miRNA-seq datasets, showing significant expression pattern differences between the isomiRs and their corresponding reference miRNAs, leading to misinterpretation of differential expression signals for both. As a case study, we generate a new dataset assessing isomiR abundance under hypoxia in human endothelial cells between the nuclear and cytosolic compartments. The results suggest that isomiRs are dramatically altered in their nuclear localization in response to hypoxia, indicating a potential non-canonical effect of the species, which would be missed without isomiR-aware analysis. Our results call for a comprehensive re-evaluation of the miRNA-seq analysis practices.

## Introduction

MicroRNAs (miRNAs) are endogenous non-coding RNAs, involved in mRNA targeting for translational inhibition or degradation (1). Initially transcribed from the genome by RNA polymerase II, their biogenesis involves a two-step cleavage process to produce mature miRNA sequences. Firstly, the primary miRNA transcript (pri-miRNA) is processed by the Drosha enzyme in the nucleus to create a precursor miRNA (pre-miRNA). Subsequently, this pre-miRNA is exported to the cytoplasm and further cleaved by the Dicer enzyme, resulting in the mature miRNA, which can guide the RNA-induced silencing complex (RISC) to target mRNAs (**Figure 1A**) (2).

**Figure 1.**
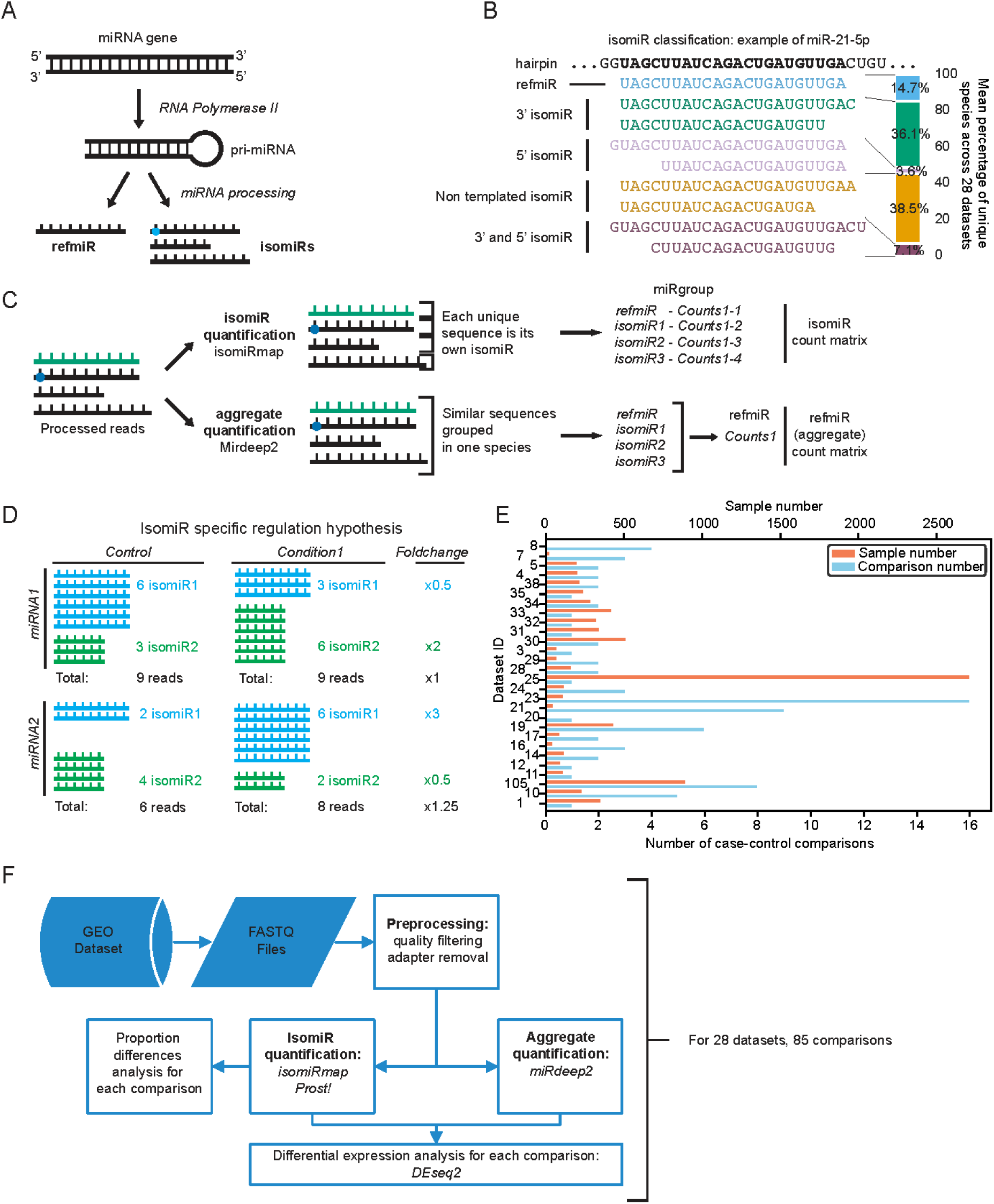
Schematic overview of the isomiR types and analysis. **A.** isomiR and miRNA biogenesis. A miRNA gene is transcribed into a primary miRNA (pri-miRNA) by the RNA Polymerase II. Various proteins, such as Drosha and Dicer, process the pri-miRNA further, cutting it into a mature miRNA sequence. In this process, alternative cleavage and additional modifications produce isoforms (isomiRs) which differ from the reference sequence (*refmiR*). **B.** isomiRs can be categorized based on their alignment to the *refmiR*: *3’ isomiRs* and *5‘ isomiRs* for 3’ and 5’ end modifications, respectively, *Non-templated isomiRs* when an addition / substitution of a nucleotide doesn’t follow the hairpin sequence pattern, and finally *3’ and 5’ isomiRs* for modifications in both 5’ and 3’ ends. On the right is depicted the mean percentage of species originating from each isomiR category, across 28 publicly available miRNAseq datasets. **C.** Overview of the computational miRNA processing with *isomiR quantification* (reads are counted independently for each isomiR sequence) and *aggregate quantification* (summing up isomiR counts from the same *miRgroup*). **D.** When isomiRs are specifically regulated in a case-control experiment, *aggregate* (in black) and *isomiR* (in blue and green) *quantifications* can yield different results. Adding up isomiR counts in *aggregate quantification* can potentially hide expression differences. **E.** Sample number (orange) and number of comparisons (blue) distributions across the 28 datasets included in the study. **F.** Overview of the processing pipeline. For each dataset, FASTQ files were obtained from GEO and preprocessed through quality filters and adapter removal. Then, isomiRmap and Prost! were used for *isomiR quantification* and miRdeep2 for *aggregate quantification*. Subsequently, DEA was performed for each comparison, and both quantification methods, using DESeq2. In addition, we ran isomiR proportion differential analysis.

The use of next-generation sequencing in miRNA research has resulted in the emergence of isomiRs – miRNA sequences that vary from their reference sequence (3). These miRNA isoforms differ from their reference sequences often just by one or two nucleotides and are a result of diverse mechanisms, such as alternative cleavage by Drosha/Dicer and non-templated nucleotide additions (4–6). The modifications from the reference sequence are the basis of current isomiR classification, which divides isomiRs into 5 categories, namely *3’ isomiRs, 5‘ isomiRs, Non-templated isomiRs* and finally *3’ and 5’ isomiRs* (**Figure 1B**) (5, 7).

Paradoxically, isomiRs have received limited attention in miRNA research, despite being a highly expressed and diverse group of RNA species (6, 8, 9). These molecules have been shown to exhibit varying expression patterns across individuals, tissues (10), cell types (11), gender (12), age brackets (13), and diseases (14–17). As a result, isomiRs are viewed as potential biomarkers in various disorders (18–20) and even as therapeutic agents (21, 22). Yet, the inclusion of isomiRs in miRNA studies remains inconsistent. This gap between their evident value and limited representation in research can be traced to two primary reasons. Firstly, the lack of standardized computational methods makes their analysis and interpretation challenging. Secondly, they are frequently viewed as merely an adjunct to the traditional miRNA analysis, rather than as a focal point of study. isomiRs could play a significant role in miRNA differential expression analysis (DEA). Neglecting them might result in missing crucial expression signals. For instance, Giuliani *et al*. (23) found that out of 133 differentially expressed (DE) isomiRs, only half showed DE in their respective reference miRNAs, when analyzed using traditional miRNA methods. Another study identified instances where isomiRs, derived from the same reference sequence, were DE in opposite directions (24). These findings not only cast doubts on the efficacy of standard methods but also emphasize the importance of understanding the role of isomiRs in DE to prevent discrepancies between computational results and biological reality before pursuing experimental validations, which usually focus on targeting or inhibiting specific sequences.

Here, we sought to compare the traditional miRNA quantification method, which merges potential isomiR signals under a primary reference, with a method that individually counts isomiR reads, providing a detailed perspective on isomiR-specific miRNA expression. For comprehensive analysis, we processed 28 publicly accessible miRNA-seq datasets, unveiling widespread and systematic DE patterns among isomiRs, with major discrepancies between the two methods of quantification, exposing the critical necessity of their incorporation in studies of miRNA responses to biological signals and stimuli.

## Summary of the terminology

**refmiR:** The reference sequence of a given miRNA from the miRNA reference sequence database, miRbase. Synonym: canonical miRNA, reference miRNA

**isomiR:** isoform of miRNA. isomiRs are defined relative to the *refmiR* as they are classified based on the reference sequence alignment (**Figure 1B**). *refmiRs* can also be considered as an isomiR class (the canonical isoform of a miRNA). The 5 categories of isomiRs are as follows: *3’ isomiR, Non-templated isomiR, 5’ isomiR, 3’ and 5’ isomiR, refmiR*.

**miRgroup:** A group of isomiR sequences that align to the same *refmiR*, and originate from the same miRNA arm. The *miRgroup* contains the *refmiR*. The *miRgroup* can also be called miRNA arm or miRNA species, but it is ambiguous as most miRNA studies do not account for isomiRs and use the term miRNA and miRNA arm as a synonym of the *refmiR* sequence.

**miRNA gene:** The gene from which all isomiRs of two miRgroups (3p and 5p arms) originate.

**isomiR quantification:** a miRNA sequencing analysis method that independently counts isomiR reads and aligns them to their respective *miRgroups*. However, this method does not aggregate the counts of isomiRs from the same *miRgroup*. Instead, it produces an isomiR-level count matrix, providing one count for each isomiR in every sample.

**aggregate quantification:** A prevalent miRNA sequencing analysis method that adds up counts from sequences aligned to the same *refmiR*. This results in counts exclusively at the *miRgroup* level, masking distinct isomiR signals. The underlying premise of this analysis is that the biological diversity of miRNAs arises solely from the *refmiR*, and isomiRs within the same *miRgroup* exhibit identical signals and functions. This method is also referred to as canonical, classical, or standard quantification/analysis. In this work, we employ miRdeep2 for this quantification, given its widespread citation in the literature.

## Materials and methods

### Cellular compartment / Hypoxia dataset

Human Umbilical Vein Endothelial Cells (HUVECs) were extracted with collagenase (0.3 mg/ml) digestion from umbilical cords obtained from the maternity ward of the Kuopio University Hospital. The collection was approved by the Research Ethics Committee of the Hospital District of Northern Savo, Kuopio, Finland. Written informed consent was obtained from the donors. The cells were cultivated in Endothelial Cell Basal Medium (Lonza) with recommended supplements (EGM SingleQuot Kit Supplements & Growth Factors, Lonza). Cells of seven donors were used in the study in separate, unpooled batches. All results were repeated on at least three donor batches. For the compartment dataset, HUVECs (passage 6) were cultured in T75 flasks in humidified CO2-incubator (0h control cells) or in a hypoxia chamber (Baker Ruskinn) with 1% O2, 5% CO2 for 7h or 24h. Cells were washed with PBS and collected by scraping to PBS+0.5% BSA. Cells were pelleted by centrifugation at 700g, +4 °C for 5 min and washed by PBS+0.5% BSA. Cell pellets were lysed with a hypotonic lysis buffer and nuclear and cytoplasmic fractions were isolated according to the protocol by Gagnon et al. (25). Total nuclear and cytoplasmic RNA were extracted using TRIzol Reagent (Thermo Fisher Scientific) and RNA was dissolved into molecular biology grade water. Total RNA was treated with DNase I (cat. EN0521, Thermo Fisher Scientific) according to manufacturer’s instructions. RNA Clean & Concentrator-5 kit (cat. R1013, Zymo Research) was used to separate both nuclear and cytoplasmic total RNA into long and small RNA containing fractions according to the manufacturer’s protocol. RNA quality was assessed using the Standard Sensitivity RNA analysis kit (cat. DNF-471-0500, Agilent Technologies) Fragment Analyzer. Nuclear and cytoplasmic fraction separation was confirmed by qPCR for tRNA (htRNA-Lys-TTT-3-4). cDNA was synthesized using RevertAid Reverse Transcriptase (Thermo Scientific) and gene-specific primer (reverse primer) and quantified using Maxima SYBR Green/ROX qPCR Master Mix (2×) (Thermo Fisher Scientific). Thermal cycling was performed using a LightCycler480 (Roche) with the following program: 10 min at 95 °C, followed by 50 cycles of 15 s at 95 °C and 60 s at 60 °C. Primers used were (sequences are 5D to 3D) Forward: GCCCGGATAGCTCAGTCG and Reverse: CGCCCGAACAGGGACTTG. The libraries were prepared using the NEBNext® Multiplex Small RNA Library Prep Set for Illumina according to the instructions. Library sequencing was performed on the NextSeq 500 platform.

### Public datasets - acquisition

All 28 datasets included in this study were obtained from GEO. They were primarily selected for the diversity of biological context and origin, in addition to the number of samples. We provide a detailed description of their GSE number, and reference paper, as well as other information (**Supplementary Table 1**). The SRA platform was used to obtain sample level information, such as biological conditions (cell type, disease, treatment) that were used to run case-control comparisons, for which we provide a detailed description (**Supplementary Table 2**). FASTQ files for each sample, of each GEO dataset were collected from SRA, using the prefetch command of *sra* tools, and *fasterq-dump*.

### miRNA-seq preprocessing

When necessary, adapter sequences were removed from the reads (in some cases, the submitted FASTQ files, available on SRA, already had their adapter removed). Adapter removal was performed using *cutadapt -a [ADAPTER SEQUENCE] -m 10 --max-n 0 -j 8* (25). Then, reads were filtered by quality, to only retain reads were 100% of nucleotides have a phred score of more than 30: fastq_quality_filter -q 30 -p 100. FASTQ files were also converted to fasta files for processing with PROST! (26).

### Aggregate quantification with miRDeep2

As a conventional miRNA-seq processing pipeline, miRDeep2 (27) was used to process the fasta files. From this pipeline, the command mapper.pl -c -j -m -s was run to collapse the reads and then quantifier.pl -p ../human_hairpin.fa -m ../hsa_mirna.fa -r -t hsa -d -j -y to align the sequences to the miRNA miRbase sequences (*refmiRs*) and their precursor. The resulting count files were then merged together. As miRDeep aligns not only to mature sequences, but also to precursors, some reads were counted twice and several rows of the same mature sequence existed in the counts table. This was corrected by removing duplicate rows.

### isomiR quantification with isomiRmap and Prost!

isomiR sequences were quantified, resulting in an isomiR count matrix using isomiRmap (7) and Prost! (26).

For isomiRmap, each sample was processed using the miRBase mapping bundle, running this command python3 IsoMiRmap.py --m MappingBundles/miRBase.

For Prost!, Version 0.7.3 was used according to the documentation of the program. The software was configured to only include sequences that had a total number of 25 reads across the two datasets, as well as a minimum length of 17 nucleotides and maximum of 25. Reads were aligned to the human genome GRCh38 and the list of mature miRNA sequences and miRNA hairpins were obtained from miRBase v22 (28). In our analysis, we used count matrices and isomiR annotations from isomiRmap as it is more commonly used and has a strict filtering process for isomiR detection. This method ensures that all species included in further analysis are confidently identified as members of the miRNA family. Unless mentioned otherwise in the figure legend and text, all results originate from isomiRmap quantification.

### Count matrix processing

For accuracy and comparability, *isomiR quantification* and *aggregate quantification* count matrices are processed the same way. The matrices are normalized to obtain counts per million (CPM), and a cutoff of 20 counts per sample on average is applied to remove lowly expressed species. For further comparisons, we only keep species for which we have signal in both *aggregate* and *isomiR quantification*.

### Differential expression and differential distribution analysis

For each condition, DESeq2 (29) was used independently on the raw, non-normalized counts of all isomiRs, including miRBase sequences, as well as on the *aggregate quantification* of miRNA counts. P-values were then adjusted for FDR through DESeq2. Using the stats.f_oneway function from the scipy package, a one-way ANOVA test was performed on the proportions of each isomiR relative to the *miRgroup*. The isomiR proportions are calculated separately for each sample. This test aimed to determine if the variance in proportions within each isomiR was associated with the sample group in each case-control comparison. The p-values were further adjusted using the Benjamini-Hochberg (30) FDR correction for all isomiRs in every case-control comparison.

### Permutation analysis

We conducted a permutation analysis to assess the accuracy of our statistical testing and significance measures. In this analysis, we randomly reassigned labels to the data in 500 iterations, for each comparison. Afterward, we applied the DEA and differential proportion analysis methods to these permuted datasets.

## Results

### Systematic isomiR analysis over 28 publicly available datasets

In this study, 28 publicly available miRNA-seq dataset were processed using two methods of miRNA quantification: the first approach followed the standard miRNA analysis method that combines counts from sequences aligning to the same *refmiR*, yielding *refmiR* level results and merging potential isomiR signals. In this study, we call this approach the *aggregate quantification* method. In the second approach, the *isomiR quantification* method, we counted isomiR reads individually and annotated *miRgroups* through alignment, offering a nuanced isomiR-specific view into miRNA expression (**Figure 1C**). The conventional approach posits that specific isomiR expression patterns can be sidelined, consolidating everything back to the primary reference sequence. This stems from the past, when isomiRs were often dismissed as mere sequencing aberrations, especially when expression studies were predominantly performed using microarrays. The underlying assumption of *aggregate quantification* is that aligning reads to reference sequences would capture the most significant information, rendering the rest redundant. However, existing research on isomiRs provides evidence that alterations on the *refmiR* sequence affects the biological function, potency, or cellular behavior of the molecule, even when the seed sequence remains intact (31–33). If isomiRs exhibit unique regulatory patterns and profiles, these nuances could introduce biases in standard miRNA-seq analyses, where the *aggregate* levels for a specific miRNA could potentially remain consistent across different conditions, while the proportions of its isomiRs vary distinctly. In such a case, even if the miRNA held significance to a biological process through its isomiRs, the *aggregate quantification* would obscure the relevance. (**Figure 1D**).

We focused on 28 miRNA-seq datasets, representing a large diversity of tissues, cell cultures, biological conditions, and diseases (**Supplementary Table S1**), as well as different library preparation methods. Overall, our analysis encompassed 85 distinct case-control comparisons, and thousands of miRNA-seq samples (**Figure 1E**). Each dataset was acquired with a standardized pipeline, preprocessed, and analyzed, counting miRNA and isomiRs signals, and performing DEA (**Figure 1F**), providing a rich resource of standardized isomiR analysis results.

### isomiR diversity and expression predominance across datasets

In most miRNA-seq datasets, we observed a broad spectrum of isomiR species, with numbers varying from 303 to 1525 unique sequences. Among these, 22 of the 28 datasets encompassed over 900 distinct species **(Figure 2A**). The distribution of isomiR types remained relatively consistent between datasets both across species diversity (**Figure S1A**) and expression levels (**Figure S1B**). *Non-templated isomiRs* were the predominantly represented type (median of 38% of species), followed by *3’ isomiRs* (36%). Notably, while *3’ isomiRs* contributed to 38% of the expression, *Non-templated isomiRs* accounted for only 15%. Less common, both in numbers and expression, were the *5’ isomiRs* and the *3’ and 5’ isomiRs*, which represented a median of 3% and 7% of species and 1% and 2% of expression, respectively. The *refmiRs* made up 15% of species at median and 42% of total miRNA expression. In summary, at the dataset scale, isomiRs represented most of the expression, with a median of 58%, and a range between 41% and 79%.

**Figure 2.**
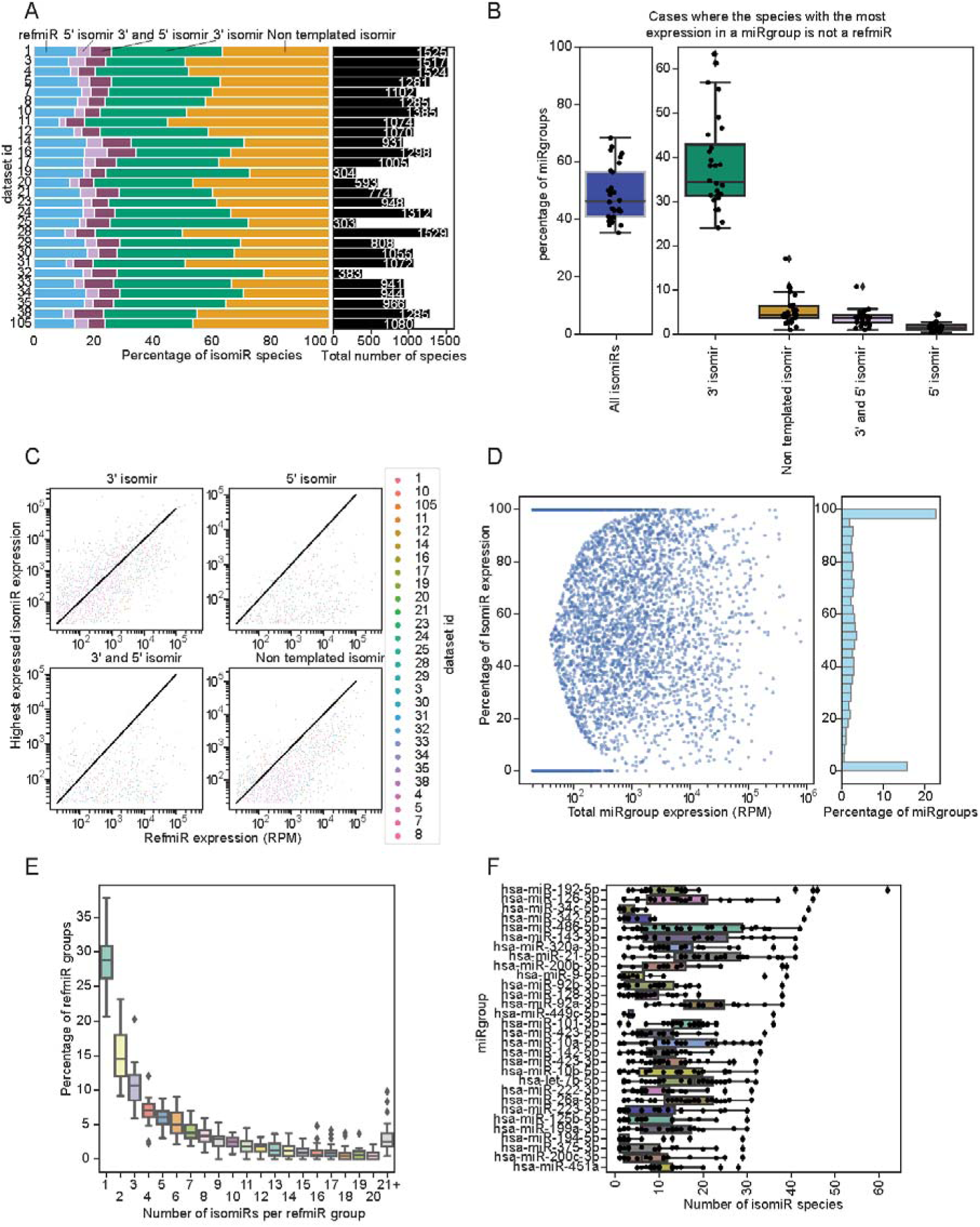
IsomiR diversity and abundance. **A.** isomiR type distribution and species count. The proportions are derived from the total number of unique species in each category. **B.** Majority isomiRs in *miRgroups*. The left panel illustrates the percentage of *miRgroups* where the dominant species (highest expression level within a *miRgroup*) is identified as an isomiR. The right panel further classifies these data points based on isomiR types. Each dot corresponds to a specific dataset, representing the percentage of *miRgroups* where the top-expressed species belongs to the isomiR category indicated on the x-axis. **C.** Comparative analysis of isomiR and *refmiR* expression levels. The scatter plots show the relationship between expression levels of various isomiR categories and their corresponding *refmiR* values. Each dot, colored based on the dataset it belongs to, represents an isomiR-*refmiR* pair, with its position determined by their respective expression magnitudes. The four plots represent the four distinct isomiR categories: *3’ isomiR*, *5’ isomiR*, *3’ and 5’ isomiR*, and *non-templated isomiR*. The black diagonal line in each plot serves as a reference, indicating points where the isomiR and *refmiR* expression levels are equivalent. **D.** Dissection of isomiR contribution to overall *miRgroup* expression. The scatter plot on the left shows a visual representation of the proportion of isomiR expression (y-axis) relative to the total *miRgroup* expression (x-axis). Each dot corresponds to a distinct *miRgroup* from various datasets. The x-axis is log-scaled. Adjacent to the scatter plot, the histogram on the right shows a frequency distribution for the percentage of *miRgroups* (y-axis) at specific isomiR expression percentage intervals (x-axis). **E**. Distribution of isomiRs within *miRgroups* across datasets, showing the variation in percentages across different datasets for each isomiR count. **F.** Variability in isomiR counts for highly expressed *miRgroups* across datasets. This graph showcases the distribution of isomiR counts for the *miRgroups* with the highest levels of expression. On the y-axis, specific *miRgroups* are listed, while the x-axis indicates the number of isomiRs found within those groups. Each data point represents the count of isomiRs for a particular *miRgroup* in a given dataset.

A significant proportion of the expression came from isomiRs, leading us to closely examine the dominant species at the *miRgroup* level. Our analysis sought to discern how frequently an isomiR emerged as the majority species over *refmiRs*. Although the underlying assumption of the *aggregate quantification* suggests that the *refmiR* should be the most expressed species in a *miRgroup*, this was not consistently observed. In fact, isomiRs served as the majority species in a notable range of cases, with instances varying from 35% to 68% of *miRgroups*, and a median of 46% (**Figure 2B**). While the *3’ isomiRs* often dominated these instances (with a median of 30% majority species), occurrences of *non-templated isomiRs, 5’ isomiRs, or 3’ and 5’ isomiRs* being the majority species were also observed (**Figure 2B**). This trend persisted across isomiR types. Although *3’ isomiRs* were clearly predominant in expression, in many instances the most expressed isomiR of any type outpaced the *refmiR* in its *miRgroup* (**Figure 2C**). Importantly, this pattern spanned across diverse expression levels (encompassing low, moderate, and high expression) indicating a widespread phenomenon.

To further assess isomiR dominance within *miRgroups* and contextualize it within the diversity of miRNA expression levels, we studied the distribution of isomiR expression across *miRgroup* abundance levels (**Figure 2D**). A consistent pattern emerged: lowly expressed *miRgroups* were either all *refmiR* or all isomiR. With higher expression level, we saw a uniform distribution of isomiRs throughout various *miRgroups*. isomiRs consistently represented a significant fraction of the expression in the higher expression levels (**Figure S1C**) and across datasets (**Figure S1D**). In 80% of *miRgroups*, isomiRs accounted for more than 20% of their expression, while 60% of *miRgroups* had more than half of their expression originating from isomiRs (**Figure S1E**). In addition, all isomiR types were represented in most expression levels (**Figure S1F**), excluding *5’ isomiRs* from levels > 20 000 RPM, and *3’ and 5’ isomiRs* from expression levels > 50 000 RPM.

Another dimension of diversity is the variety of isomiRs within each *miRgroup*. Upon examination, the distribution of distinct isomiRs per *miRgroup* remained stable across datasets (**Figure 2E**). While the majority of *miRgroups* typically featured three or fewer isomiRs (**Figure S1G**), a significant portion of *miRgroups* harbored over 20 isomiRs, (**Figure S1H**). This suggests that the majority of isomiRs stem from a limited subset of *miRgroups*. Delving deeper into specific *miRgroups* revealed great diversity in isomiR counts between datasets (**Figure 2F**). For instance, miR-21-5p and miR-486-5p exhibited a broad spectrum, ranging from three to over 40 isomiRs, with variations evident throughout the datasets. This indicates that isomiR distribution can significantly differ across various biological settings. The observation led us to further investigate isomiR DE patterns across distinct biological conditions.

### DE discrepancies arising from isomiR expression

After confirming the significance of isomiRs in both diversity and expression levels, we explored their distribution shifts in response to specific biological conditions. For this purpose, DEA was employed to evaluate expression variations in case/control comparisons. Each comparison yielded two distinct sets of miRNA outcomes: a DE result at the *miRgroup* level from the *aggregate quantification* and at the isomiR level from the *isomiR quantification* (**Figure 3A**). A preliminary observation revealed a strong correlation between the proportion of DE species at the *isomiR* DE level and at the *aggregate* DE level (**Figure 3B**, **Figure S2A**). Our primary focus was on elucidating the putative comparative advantage of *isomiR quantification* over *aggregate quantification* for DEA. To this end, we focused on comparisons and datasets that significantly affected the expression of miRNAs. As such, we established a 20% threshold: only those comparisons where at least 20% of miRNAs demonstrated significant *aggregate* DE were considered (**Figure 3B, Figure S2A**).

**Figure 3.**
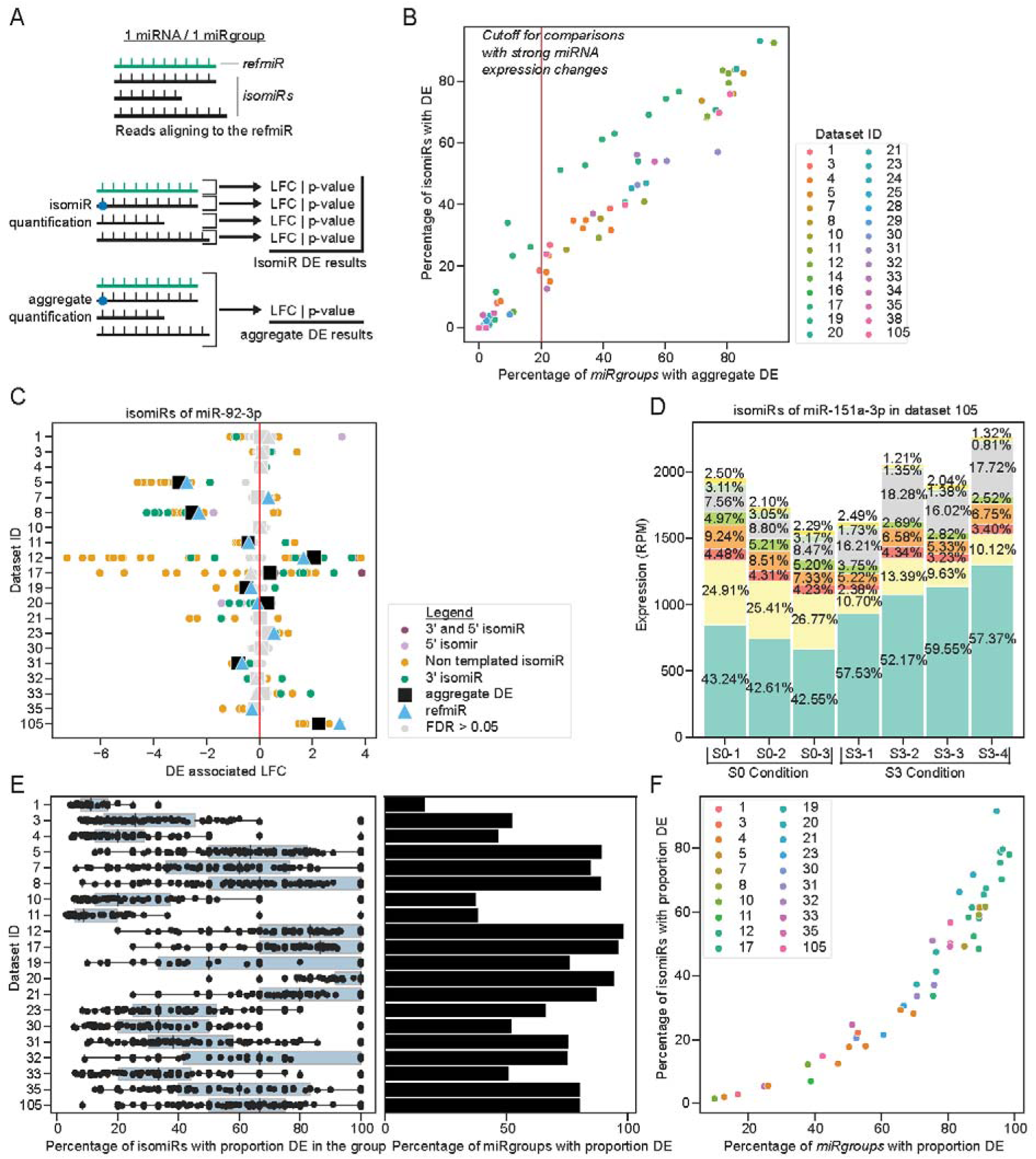
IsomiR differential expression patterns highlight complex and specific distribution. **A.** DE with *isomiR quantification* and *aggregate quantification* count matrices. **B.** Overview of miRNA signals across comparisons DE using *aggregate* (x-axis) and *isomiR* (y-axis) *quantifications*, presented as the percentage of species showing significant DE. A threshold of 20% in *aggregate quantification* is used to focus on comparisons with pronounced miRNA signals. **C.** DEA of miR-92-3p across top comparisons in each dataset. Displayed are results for *aggregate quantification* (black square) and *isomiR quantification* (dots, color-coded by isomiR type), which includes the *refmiR* (blue triangle). The x-axis shows the LFC for each comparison. Significant cases are color-coded, whereas non-significant results (FDR > 0.05) are presented in gray. **D.** Proportional changes in isomiR expression for miR-151a-3p in dataset 105. The bars represent individual samples under conditions S0 and S3. Different colors within the bars correspond to distinct isomiRs. The percentages indicate the expression contribution of each isomiR relative to the total *miRgroup* expression for that particular sample. **E.** Distribution of *miRgroup* proportion changes across datasets. On the right, the percentage of *miRgroups* in each dataset that contain at least one isomiR showing a significant proportion change is presented. On the left, within those *miRgroups*, the distribution of isomiRs displaying significant proportion changes is depicted. Each line represents the top comparison for the respective dataset. **F.** Scatter plot displaying the relationship between the percentage of isomiRs (y-axis) and *miRgroups* (x-axis) with significant distribution differences. Each dot represents a distinct comparison, differentiated by dataset color. A *miRgroup* is marked as having a distribution change if at least one of its isomiRs exhibits such a change, significantly.

The main question that arises when seeking to assess the impact of isomiRs in DE results is, if the DE signal of isomiRs differs within the same *miRgroup*.

A way to answer this is to observe the Log2FoldChange (LFC) of the isomiRs and *aggregate quantification* within the same *miRgroup*. For example, for miR-92-3p, which was selected among *miRgroups* displaying high levels of isomiR DE, a wide range of LFC was observed (**Figure 3C**) and many isomiRs even showed opposite LFCs compared to *aggregate quantification*.

However, LFC alone is not ideal for capturing isomiR-specific signals as it will depend on both the general *miRgroup*-associated changes (responding to miRNA gene expression changes), and the isomiR-specific regulation. To explore the effects of biological changes specifically on isomiR distribution, and independently of the *miRgroup* level changes, we investigated isomiR expression proportions within the same *miRgroup.* We defined this proportion as the percentage of reads originating from a specific isomiR across all isomiRs of the same *miRgroup*, in each miRNA-seq sample. If an isomiR proportion significantly changed between conditions, that would constitute a change of expression / regulation, specific to the isomiR. This is the case for example in the senescence dataset (ID 105) for miR-151a-3p, that showed a significant increase in proportion for some isomiRs, and decrease for others, between the conditions S0 and S3 (**Figure 3D**). This contrasts with random distribution of isomiRs where we would expect random or stable proportions within a *miRgroup* across conditions.

Expanding the isomiR proportion difference analysis for all datasets and *miRgroups,* we found a high percentage of isomiRs with statistically significant proportion changes across all datasets (**Figure 3E**). Over all comparisons, the percentage of isomiRs displaying significant proportion changes ranged from 9.8% to 98.5%, with a median of 76.5% (**Figure S2B**). When selecting one comparison for each dataset (selecting the one with the most overall miRNA DE signal, **Figure S2C**), values were a minimum of 17%, maximum of 98.5% and median of 75.9% (**Figure S2D**). In *miRgroups* with at least one variably distributed isomiR, we observed a high frequency of isomiRs with significant proportion differences (**Figure 3E**). Across all isomiRs, we saw a systematic and pronounced trend, with a vast majority of cases exhibiting such differences (majority of isomiRs, and majority of *miRgroups*) (**Figure 3F**). Taken together, our data underscores that variations in isomiR proportions are not only widespread but also consistent across the datasets and comparisons analyzed, suggesting specific regulation and response to biological signals.

After determining the presence of isomiR-specific signals in differential distribution, we then asked: to what degree does this lead to inconsistencies or missed signals when relying solely on *aggregate quantification* DE? Using miR-92-3p as an example (**Figure 3C**), we juxtaposed *isomiR* DE against its corresponding *aggregate* DE, analyzing each dataset. We categorized *isomiR* DE results into distinct groups based on their relationship to *aggregate quantification* DE (**Figure 4A, Figure S2E**):

- *Opposite DE:* Both *aggregate* and *isomiR* signals indicate significant DE, but they diverge in terms of LFC direction.
- *Same DE:* Both signals exhibit significant DE, with the LFC pointing in the same direction.
- *Only isomiR DE:* The *isomiR* displays significant DE, while the corresponding *aggregate* signal does not.
- *Only aggregate DE:* The *isomiR* does not exhibit significant DE, but its related *miRgroup* does at the *aggregate* level.

**Figure 4.**
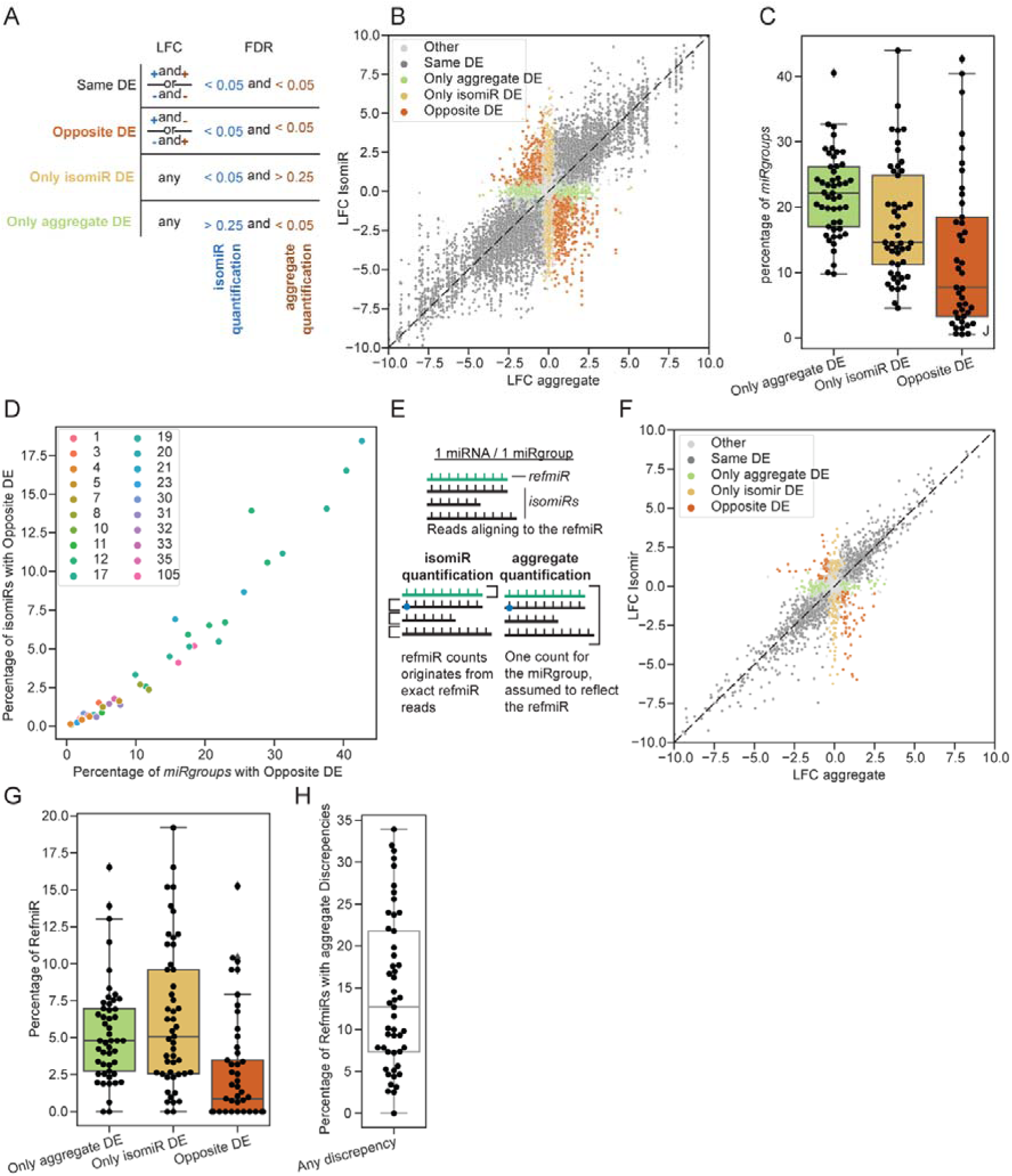
Systematic differential expression patterns of isomiRs highlights discrepancies with *aggregate quantification*. **A.** Categorization schema comparing DE signals from *aggregate quantification* and *isomiR quantification*. Each isomiR is categorized based on its log fold change (LFC) and false discovery rate (FDR) values in *isomiR quantification* DE, as well as the LFC and FDR values of its corresponding *miRgroup* in *aggregate quantification* DE. **B**. *isomiR* vs *aggregate* associated LFC for all comparisons, colored by classification. The scatter plot illustrates the relationship between the *aggregate quantification DE* LFC l(LFC aggregate) and *isomiR*-specific LFC (LFC isomiR). Points are color-coded based on the DE classification: *Other* (gray), *Same DE* (black), Only *aggregate DE* (green), Only *isomiR DE* (yellow), and *Opposite DE* (orange). The dashed line indicates where the LFC values for *isomiR* and *aggregate* are equal. **C**. Distribution of the number of *miRgroups* that show discrepancies between *aggregate* and *isomiR* DE. The box plots display the percentage of *miRgroups* for three categories: Only *aggregate* DE (green), Only *isomiR DE* (yellow), and *Opposite DE* (orange). Each point represents a specific *miRgroup*, and the spread of the points shows the variation within each category, across datasets. **D**. Percentage of isomiRs (y axis) and *miRgroups* (x axis) with *Opposite DE* classification, across all comparisons. The scatter plot displays the relationship between the percentage of *miRgroups* with *Opposite DE* and the percentage of *isomiRs* with *Opposite DE.* Each point is color-coded based on a specific dataset number, indicating the comparison for that dataset. A *miRgroup* is considered to have *Opposite DE* if at least one of its isomiR is classified as such. **E**. *isomiR* vs *aggregate quantification* and its significance for the *refmiR* sequence. The *aggregate quantification* method combines all isomiRs of a *miRgroup*, interpreting them as aligned or corresponding to the *refmiR* sequence. In contrast, the *isomiR quantification* treats the *refmiR* as its own species within the *miRgroup*, allowing for unique expression and distribution measurements. **F**. Scatter plot showing the relationship between LFC values sourced from the *refmiR* (vertical axis) and its corresponding *aggregate* signal (horizontal axis). **G**. Box plot illustrating the distribution of percentages for *refmiRs* that deviate in DE from *aggregate* signals, grouped by specific discrepancy type. **H**. Representation of all discrepancies between *refmiR* and *aggregate* signals, presented as a percentage.

Among these, *Opposite DE* and *Only isomiR DE* are of particular interest as they indicate potential inconsistencies or signals overlooked with *aggregate quantification* DE. In fact, nearly every dataset revealed at least one isomiR from these two categories for miR-92-3p (**Figure S2E**). Expanding our analysis to encompass all *miRgroups* and comparisons revealed marked variance in isomiR LFC, within isomiRs of the same *miRgroup* (**Figure 4B**). While most isomiRs overall agreed with their corresponding *aggregate* DE outcomes, numerous instances of *Opposite DE* and *Only isomiR DE* emerged throughout datasets. They constituted a significant proportion of *miRgroups* (**Figure 4C, Figure S3A**) and isomiRs (**Figure S3B-C**). When evaluating the *miRgroups* with at least one *Opposite DE* species (**Figure 4D**), it became evident that, even if this signal is not as dominant as in proportion differences, it is unquestionably widespread. In some instances, up to 40% of *miRgroups* had at least one *Opposite DE* isomiR. This underscores major inconsistencies and potential missed signals from isomiRs when relying solely on *aggregate quantification*. Such patterns emerge consistently in notable numbers across datasets and comparisons.

In *isomiR quantification*, *refmiRs* were incorporated into the expression matrix just like any other isomiR. Specifically, only reads that perfectly aligned with the *refmiR* sequence were participating in the *refmiR* count (**Figure 4E**). Contrastingly, *aggregate quantification* amassed counts from isomiRs within the same *miRgroup*, attributing the resulting count to the *refmiR* and effectively sidelining the isomiRs. This produced two unique counts for the *refmiR*. Discrepancies between the specific *refmiR* DE signal and the overall *aggregate quantification* DE were of paramount interest. Our analysis revealed that significant deviations occured, with marked LFC differences (**Figure 4F**). Instances of *Opposite DE*, *only aggregate DE*, and *only refmiR DE* were prevalent across the majority of datasets and comparisons, covering a broad spectrum of expression levels, inclusive of highly expressed miRs (**Figure S3D**). This prevalent pattern impacted up to 30% of species (**Figure 4G-H**), with a median value of 12% across all comparisons. Among these, *Opposite DE* cases were particularly interesting (reaching up to 15% of all species), but all discrepancies were highly relevant. The *Only aggregate DE* cases may result in false positives, if the *miRgroup* is DE at the *aggregate* level, but the actual *refmiR* is not, when considering *isomiR quantification*. On the flip side, scenarios where the *refmiR* show significant DE through *isomiR quantification*, but not in *aggregate* (*only refmiR DE*), suggest false negatives.

IsomiRs that possess altered seed sequences are of special interest when examining the influence of *isomiR quantification* on DE. A shift in the seed sequence would considerably increase the likelihood that an isomiR had a distinct cellular role compared to the *refmiR*, given that a canonical miRNA target is predominantly determined by its seed sequence. Interestingly, the proportion of isomiRs showing significant variances was strikingly consistent, whether their seed sequence differed from their *refmiR* or not (**Figure S4A**). In most datasets and comparisons, there were at least 20 isomiRs with seed sequence alterations that also displayed differential distribution at the proportional level (**Figure S4B**). Moreover, discrepancies with *aggregate quantification*, such as *opposite DE* or *only isomiR DE* cases were observed at similar levels among species with or without seed changes (**Figure S3C**). This consistency was also apparent across various isomiR types (**Figure S3D**). Such findings underscore the significance of these discrepancies. The instances were not anomalies; they were prevalent and systematic across multiple datasets and comparisons, most of which presented a considerable number of discrepancies among species with seed alterations (**Figure S4E-F**).

We then sought to validate the statistical significance of the observed differences in isomiR expression patterns, as DESeq may not fully accommodate the unique distribution characteristics of isomiR expression in DEA. To this end, we implemented a permutation analysis that we performed in each case-control comparison, for all datasets. We randomly reassigned the sample case/control labels, 500 times, reapplying the DEA and differential proportion analysis methods to these permuted datasets. The results, as depicted in **Figures S5A-D**, demonstrated that permutations yielded significantly fewer species with significant proportion changes and significant DE compared to actual data. Upon closer examination, the counts of differentially expressed and distributed isomiRs in each actual comparison were consistently higher than the maximum count observed in any permutation, where most observed counts were near zero. In most comparisons, fewer than 10 DE isomiRs were detected in more than 98% of the comparisons. These results strongly support the validity of our method and substantiates the statistical significance of our findings.

In addition, we investigated potential biases affecting our findings, such as the library preparation method (**Figure S6**) and isomiR detection methods (**Figure S7**). Our analysis suggests that while different library preparation methods might show varying isomiR distribution patterns, discrepancies in both *aggregate* and *isomiR quantification*, including those involving *refmiRs*, were consistently observed across all methods (**Figure S6**). Additionally, our comparison of *Prost!* (26) and *isomiRmap* (7) revealed a uniform level of these discrepancies (**Figure S7**), reinforcing the reliability of our findings.

Our findings attest to the nuanced condition-specific signals arising from the isomiRs. This signal is consistently evident across datasets and comparisons, sometimes in astonishing quantities. If analysis is restricted to the *aggregate quantification*, significant signals can be missed, leading to potential discrepancies that affect *refmiRs* themselves. This raises concerns over the conclusions of the *aggregate quantification* studies that impact subsequent experimental validation studies and potential structural analyses.

### isomiR abundance varies between cellular compartments in response to hypoxia

Recent studies have elucidated non-canonical miRNA function both in the cytosol and the nucleus in cardiovascular disease context (34–36) prompting the need to better understand the distribution patterns of miRNAs and isomiRs between the cytosol and nucleus in response to disease-relevant stimulus. To this end, we generated a dataset in HUVECs under hypoxia (7h, 24h) and normoxia (0h). We separated cytoplasms from nuclei for each sample and extracted RNA from both fractions independently (**Figure 5A)**. We focused the analysis on three specific aspects: First, we investigated the significance of *isomiR quantification* with the same analytical methods as before, constituting a validation of our previous findings in the publicly available datasets. Then, we studied isomiR expression patterns across different cellular compartments—an area that has not been previously explored. Finally, our analysis uncovered how hypoxia influenced the expression patterns of isomiRs between the compartments.

**Figure 5.**
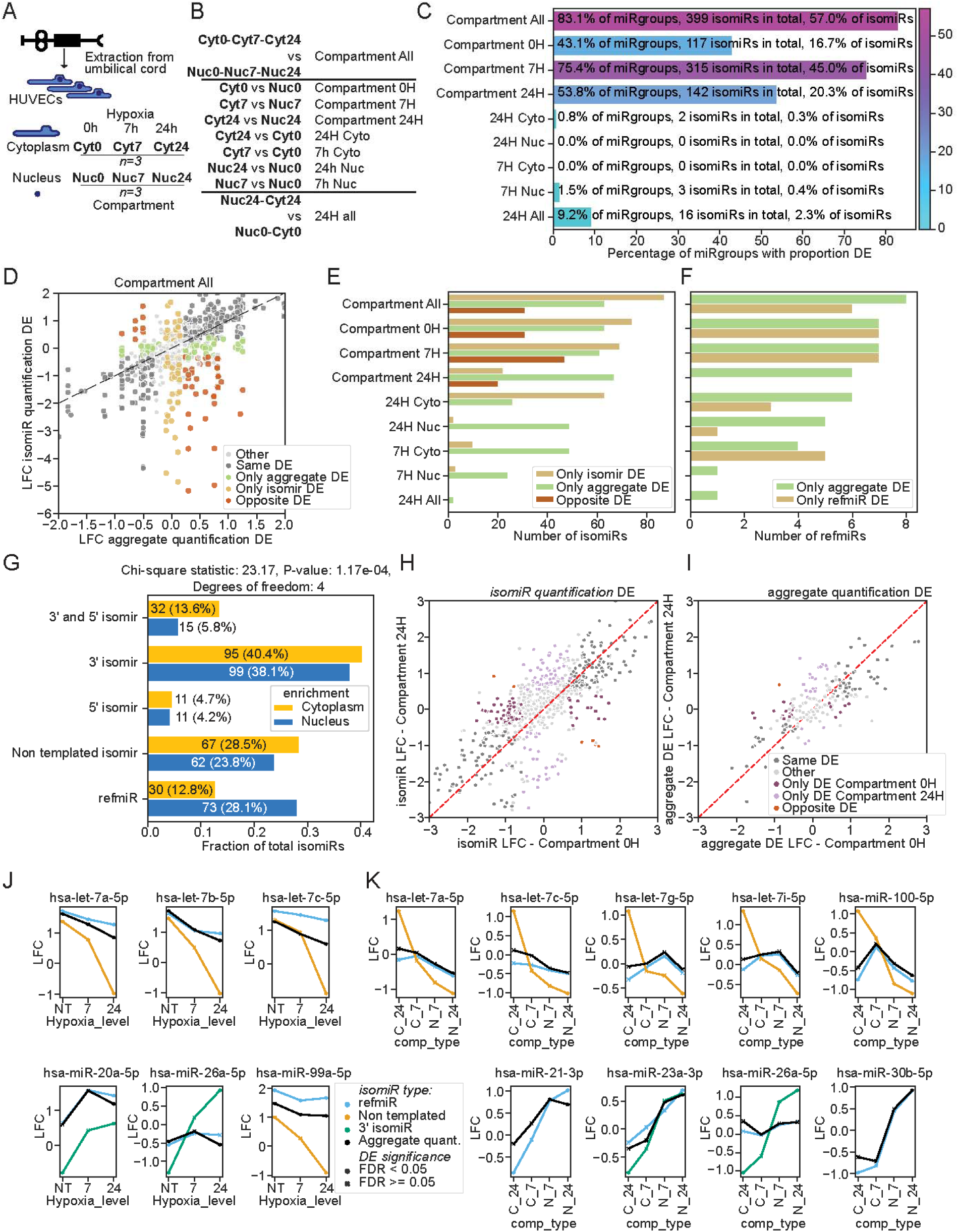
Investigating changes in isomiR gene expression under hypoxic conditions within the nuclear and cytoplasmic compartments, in HUVECs. **A**. Schematic representation of the experimental workflow. Human Umbilical Vein Endothelial Cells (HUVECs) are obtained from the umbilical cord. Following extraction, these cells are exposed to hypoxia at distinct intervals: 0 hours (0h), 7 hours (7h), and 24 hours (24h). For each time point, both the cytoplasmic (Cyto) and nuclear (Nuc) components are isolated. Subsequent to the separation, RNA from these compartments is extracted for sequencing. Each condition is tested in triplicate (N=3). **B**. Comprehensive list of comparisons made during the study. The comparisons are classified into two main categories. *Compartment Comparisons*: These are assessments made to understand the nuclear and cytoplasmic enrichments under different conditions. For example, *Compartment 0h* observes the differences between the cytoplasm and nucleus at the 0h hypoxia time point (normoxia). *Hypoxia-induced Expression Comparisons*: These are evaluations designed to discern the expression changes occurring due to hypoxia, either within the cytoplasmic or nuclear compartment. For example, *24h Cyto* examines the change in expression within the cytoplasm from the start (0h) to 24h post hypoxia initiation. **C.** Graph illustrating the proportion changes across various comparisons (y-axis) and the percentages of *miRgroups* where at least one isomiR exhibited notable proportion variations (x-axis). The color intensity of the bars corresponds to the percentage of isomiRs within that *miRgroup* with significant proportion differences. D. *IsomiR* vs *aggregate* associated LFC for all comparisons, colored by classification. The dashed line indicates where the LFC values for *isomiR* and *aggregate* are equal. **E.** Distribution of the number of isomiRs that show discrepancies between *aggregate* and *isomiR* DE, across comparisons. **F.** Distribution of the number of *refmiRs* that show discrepancies between *aggregate* and *isomiR* DE, across comparisons. **G.** Number and percentage of isomiRs that are enriched in the cytoplasm (blue) or nucleus (orange), grouped by isomiR type (y axis). Chi-square statistics is run to test for independence of distribution of isomiR type and enriched compartment within significantly enriched isomiR species. **H-I.** Comparison of nuclear enrichment in hypoxic (y axis) and normoxic (x axis) conditions, across *isomiR* (**H**) and *aggregate quantification* (**I**). Discrepancies between hypoxic and normoxic compartment enrichments are colored. **J-K.** Cases where nuclear enrichment reverses with hypoxia (**J**) and where hypoxia DE direction reverses between compartments (**K**). Dots have adjusted p-value < 0.05, crosses do not. Color refers to the signal origin, isomiR (by type, including *refmiR*), or *aggregate quantification* (in black).

We examined nine comparisons in this dataset (**Figure 5B**), either exploring compartment enrichment between different timepoints or hypoxia-associated changes between the nucleus and cytoplasm. IsomiR proportion distribution changed significantly with compartment, with up to 80% of *miRgroups* containing at least one isomiR that was significantly differentially distributed (**Figure 5C**). Much lower numbers were observed in hypoxia. These results suggest major differential isomiR distribution between the compartments, which we confirm with DEA at the *isomiR quantification* level **(Figure 5D)**. Comparison of isomiR DE results with *aggregate quantification* DE results systematically yielded discrepancies, more prominent in the compartment enrichment comparisons (**Figure 5E**). Interestingly, such discrepancies also concerned several *refmiRs* for most comparisons (**Figure 5F**), and for the hypoxia-related signals, albeit in lower numbers. A compartment-wise breakdown revealed that while *refmiRs* were primarily localized in the nucleus, *3’* and *5’ isomiRs* had a stronger presence towards the cytoplasm (**Figure 5G**). We statistically confirmed the difference of distribution in isomiR types observed between the nuclear and cytoplasmic enrichments.

We further explored the interplay between compartmental expression changes across the different hypoxia timepoints. The first observation from the *isomiR quantification* DE was that in many instances, a specific isomiR demonstrated different compartmental enrichments under normoxic and hypoxic conditions, both being statistically significant (**Figure 5H**). We also encountered situations where an isomiR displayed significant enrichment in one state but not in the other. Compared to *aggregate quantification* DE, these disparities were more pronounced for the isomiRs (**Figure 5I**), highlighting the added value of the *isomiR quantification*.

Instances where an isomiR was found to be concentrated in the nucleus under hypoxic conditions but not in normoxia (and vice versa) were of particular interest, as they suggest a potential role of hypoxia in modulating isomiR localization. We observed six such cases, for which neither the *aggregate* signal nor the associated *refmiR* showed a similar pattern (**Figure 5J**). Yet, for these isomiRs, the enrichment at 7 hours of hypoxia consistently fell between that of normoxia and 24 hours of hypoxia, suggesting progression over time in response to stimulus.

After identifying isomiRs with compartmental expression changes associated to hypoxia stimulation (**Figure 5J**), we sought to identify cases where the response to hypoxia, i.e. direction of DE between hypoxic and normoxic conditions, would be different in the nucleus compared to the cytoplasm (**Figure 5K**). We observed nine such cases, which also followed a pattern, where changes observed at 7 hours in comparison to no treatment were generally more moderate than those at 24 hours. In some cases, such as the isomiRs of 26a-5p, 100-5p, and 23a-3p, the effects at 7 hours were strikingly similar to those at 24 hours, which suggested a consistent compartment enrichment under hypoxia for these isomiRs. Collectively, our findings point to specific regulation of isomiRs under hypoxia, which results in compartment-specific differences.

Taken together, the results of our compartment dataset aligned with the findings from the public datasets and unveiled an interesting role for hypoxia in isomiR regulation that affects the cellular localization of the isomiRs. These results underscore the significance of *isomiR quantification* in unraveling their intricate biological function and poses a question of the role isomiRs play in the nucleus.

## Discussion

In this study, we present evidence advocating for the utilization of systematic *isomiR quantification*, suggesting its potential either as an enhancement or an alternative to traditional *aggregate quantification*. We pinpoint several shortcomings of *aggregate quantification*, emphasizing its tendency to yield results that can be both incomplete and misleading. Our data confirms the ubiquitous presence of isomiRs, highlighting their sequence-specific patterns and remarkable diversity. Notably, their DE often presents a contrasting picture compared to the *aggregate quantification*, showcasing in many instances an enhanced sensitivity. One of the pronounced findings was the consistent discrepancy between signals from the *aggregate quantification* and those arising directly from the *refmiR* sequence.

The robust nature of our results stems from our comprehensive approach, analyzing 28 datasets and conducting over 100 comparative evaluations. The credibility of these outcomes is further reinforced by the dataset we generated for this study. This expansive analysis allowed for a deeper dive into the intricate dynamics in various biological contexts.

Our study offers biological and methodological insights but also introduces a valuable isomiR resource, analyzing publicly available datasets that were previously unexplored at the isomiR level. Moreover, the compartment-hypoxia dataset demonstrates that a disease-relevant stimulus can cause changes in isomiR and miRNA expressions between nuclear and cytosolic compartments, with hypoxia greatly influencing their levels within and between compartments. These findings hint at hypoxia-induced roles for the molecules in both the nucleus and cytoplasm, driven by distinct regulatory pathways. This is of particular interest regarding the non-canonical role of miRNAs in the nucleus.

As canonical function of miRNA species involves the target hybridization through the seed sequence, 5’ modifications can easily modify the target selection and the function of the isomiR compared to its *refmiR*, as is supported by many studies (45–48). Nevertheless, other isomiR types, especially the highly diverse group of *3’ isomiR*, can also be of functional importance, as the 3’ end composition affects Argonaute 2 affinity for canonical miRNA function (31), and 3’uridilated isomiRs can target different mRNA molecules than their references, through tail-U-mediated repression (TUMR) (49), confirmed in specific examples (50, 51). In addition, many other non-canonical functions of miRNAs have been described, in different disease contexts, such as cancer (52) and cardiovascular disease (36), with a specific interest in gene regulatory functions in the nucleus (53). Many of these non-canonical functions involve nucleotides outside of the miRNA seed sequence, which, combined with our extensive nuclear enrichment analysis, points to a potentially high relevance of isomiRs in such functions. To confirm and explore the importance of the presented results, more studies on isomiR function are called for. Especially isomiR-specific target prediction softwares or databases would be of high interest, as most target prediction algorithms are suited for mature *refmiRs* only. Such tools could use both prediction algorithms and target discovery experiments to infer isomiR function, as current studies support both non-seed target recognition (54), and non-canonical target-gene regulation (55).

During *isomiR quantification*, several challenges and limitations arose. Given that the sequences under examination may differ as little as by only one or two nucleotides, it became essential to rely on exact sequence counts rather than alignment-based bins. This specificity could render the analysis susceptible to pitfalls like sequencing errors. However, the consistency in sequence diversity across our datasets, complemented by stringent cutoff thresholds, allowed us to mitigate the potential impact of these errors. Another challenge came from the bias introduced during library preparation, as evidenced in previous studies (37–39). These studies point to differences in the quantity and nature of isomiRs discovered between library preparation methods. Although our research identified some patterns of isomiR distribution attributed to library preparation methods, these influences were limited, as discrepancies between *aggregate* and *isomiR quantification*, including discrepancies concerning *refmiRs*, were seen in datasets across different library preparation methods. While further research avenues, like deploying paired-end sequencing (40) or randomized-end adapter protocols (39), could offer more accurate results, we found no indications that the existing biases undermined the validity of our DE and distribution findings. This is consistent with observations that library biases contribute to only a minor fraction of observed variance, with Gómez-Martín *et al.* (39) determining that library preparation artifacts accounted for only 5% of miRNA read variations.

Computational analysis of isomiRs is challenging, primarily due to the absence of a universally accepted standard for their identification. The criteria for isomiR alignment and selection remain ambiguous, and the decision on expression cutoff is often subjective. Some analytical tools, like Seqbuster (41, 42), are restrictive, permitting only specific isomiR modifications with a defined range of nucleotide changes, and while certain methodologies employ whole genome alignment (26), others rely on *refmiR* based alignments (43). Given that isomiR signals arise from both miRNA gene expression and isomiR biogenesis, there is a demand for methods integrating both aspects. Past evaluations of isomiR expression responses predominantly utilized DEA without adjusting for broad miRNA expression shifts (12, 23, 44). In contrast, our approach offers a thorough analytical framework, encompassing DE both from *aggregate quantification* and *isomiR* quantification (including *refmiR* specific insights), coupled with isomiR proportion changes. This approach extracted the maximum informational yield from miRNA sequencing. The robustness of our results was further strengthened by successful validation of statistical significance with permutation analysis, and comparison of distinct isomiR identification methods, namely Prost! (26) and IsomiRmap (7). Still, discernible need remains for refining and enhancing DEA tools tailored specifically for isomiRs.

From a broader perspective, our study challenges the prevailing approach in miRNA expression studies: the sole use of the *aggregate quantification*. This method predominantly centers on the *refmiR*, summing together all aligned reads as a singular expression count. Such an approach blurs the distinction between the *miRgroup*-level signal and the signal from *refmiR*, which represents just one sequence variant within the entire variety of isomiRs for that *miRgroup*. Our findings starkly contest this amalgamation, as we consistently identified discrepancies between the cumulative signal derived from *aggregate quantification* and the distinct signal of the *refmiR* from *isomiR quantification*. This indicates that not only crucial *isomiR* signals might be overlooked, but that *aggregate quantification* can also yield misleading outcomes, when interpreted as stemming from the *refmiR*. Given the dependence of many downstream experimental studies on miRNA-seq, pinpointing the precise sequences responsible for the signal is highly important. Even a single nucleotide alteration may carry significant implications for structural biology, induced expression, or other functions that act through yet unidentified non-canonical pathways. This observation suggests that a re-evaluation of *refmiR*-based alignment techniques may be necessary, along with a broader reconsideration of the *refmiR*-centric perspective of miRNAs and highlights the potential limitations of miRbase (28), which offers only a single reference sequence per *miRgroup*—the *refmiR*. Instead, databases that account for isomiRs, like isomiRdb (56), the largest to date, could offer a valuable alternative for referencing miRNA sequences and resolving the ambiguity between *miRgroup* and *refmiR*.

In conclusion, our results provide an updated framework for miRNA analysis that considers isomiR expression dynamics. We show that ignoring isomiRs could result in missing most of the species diversity, omitting a great amount of DE and distribution signal, and misrepresenting the reality by summing up reads from various sequences that display different or even opposing expression signals. Instead of the widely used *aggregate quantification*, we advocate for general inclusion of isomiRs in all miRNA sequencing analyses, to fully capture all relevant sequence information.

## Data availability

The associated count matrices (from aggregate and isomiR quantification), in addition to the DEA result for each case control comparison for the 28 miRNA-seq datasets presented here will be made available upon publication.

The newly generated hypoxia-compartment miRNA-seq dataset datasets presented here will be made available upon publication.

## Acknowledgements

We are grateful to Tuula Salonen for excellent technical assistance in library preparation. We acknowledge the University of Eastern Finland (UEF) Genome Center and Biocenter Finland for infrastructure support. We acknowledge Dr. Carles A. Boix and Pierre R. Moreau for their discussion and insight in this study.

