## Supplemental Figures for "The critical role of isomiRs in accurate differential expression analysis of miRNA-seq data"

### Supplementary Figures

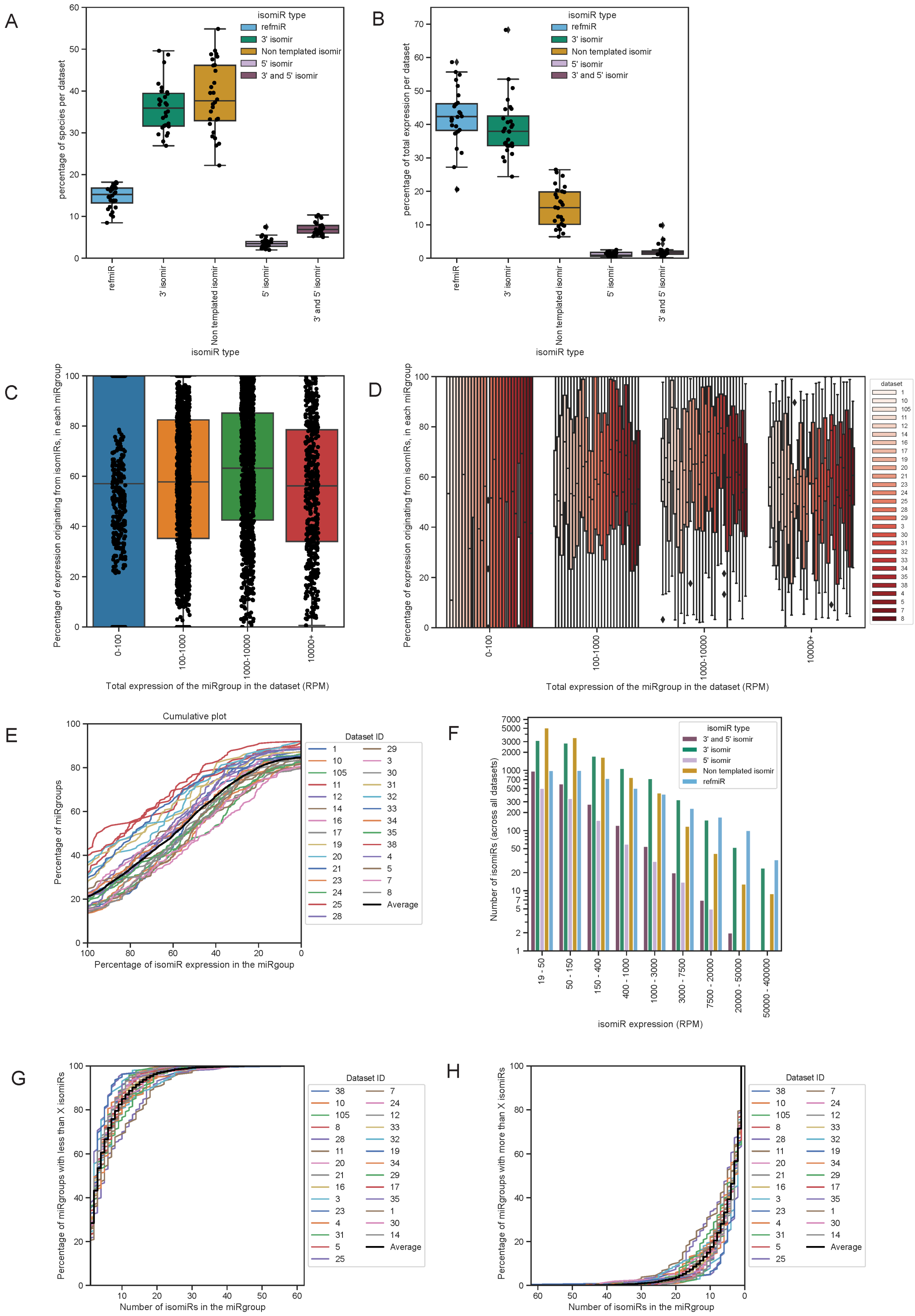

**Figure S1. Distribution and abundance of various isomiR types across datasets. A.** Distribution of different isomiR types based on the percentage of unique sequences, or species, within each dataset. Each dot on the boxplot represents a distinct dataset. The y-axis indicates the percentage of unique isomiR species present in the dataset, while the x-axis enumerates the various isomiR types. **B**. Distribution of isomiR types presented in terms of their expression levels, measured as the percentage of total reads. isomiR type diversity for each dataset, in terms of its percentage of total expression, is represented as a boxplot with each dot corresponding to a dataset. The y-axis denotes the percentage of total expression for each isomiR type, while the x-axis details the different isomiR types. **C-D**. Display of isomiR-derived expression percentages across *miRgroups*, grouped by expression levels (in RPM). Each dot symbolizes a *miRgroup* from an individual dataset, with the y-axis indicating the isomiR-based expression percentage (excluding *refmiRs*). In panel **D**, the distributions are further highlighted with boxplots, each color-coded by dataset. **E**. Cumulative distribution showing the percentage of isomiR expression within *miRgroups* for various datasets. The plot illustrates the proportion of *miRgroups* in each dataset where at least 'x' percent of their total expression stems from isomiRs, with 'x' being depicted on the x-axis. **F**. Distribution of isomiR expression levels categorized by type. The bar chart represents the count of isomiRs from all datasets, differentiated by color according to isomiR type, and grouped within specified expression ranges measured in RPM. **G-H**. Cumulative representation of isomiR count per *miRgroup*. The x-axis illustrates the total number of isomiRs within each *miRgroup*. The y-axis indicates the percentage of *miRgroups* with fewer (**G**) or greater (**H**) than 'x' isomiRs, where 'x' is specified on the x-axis.

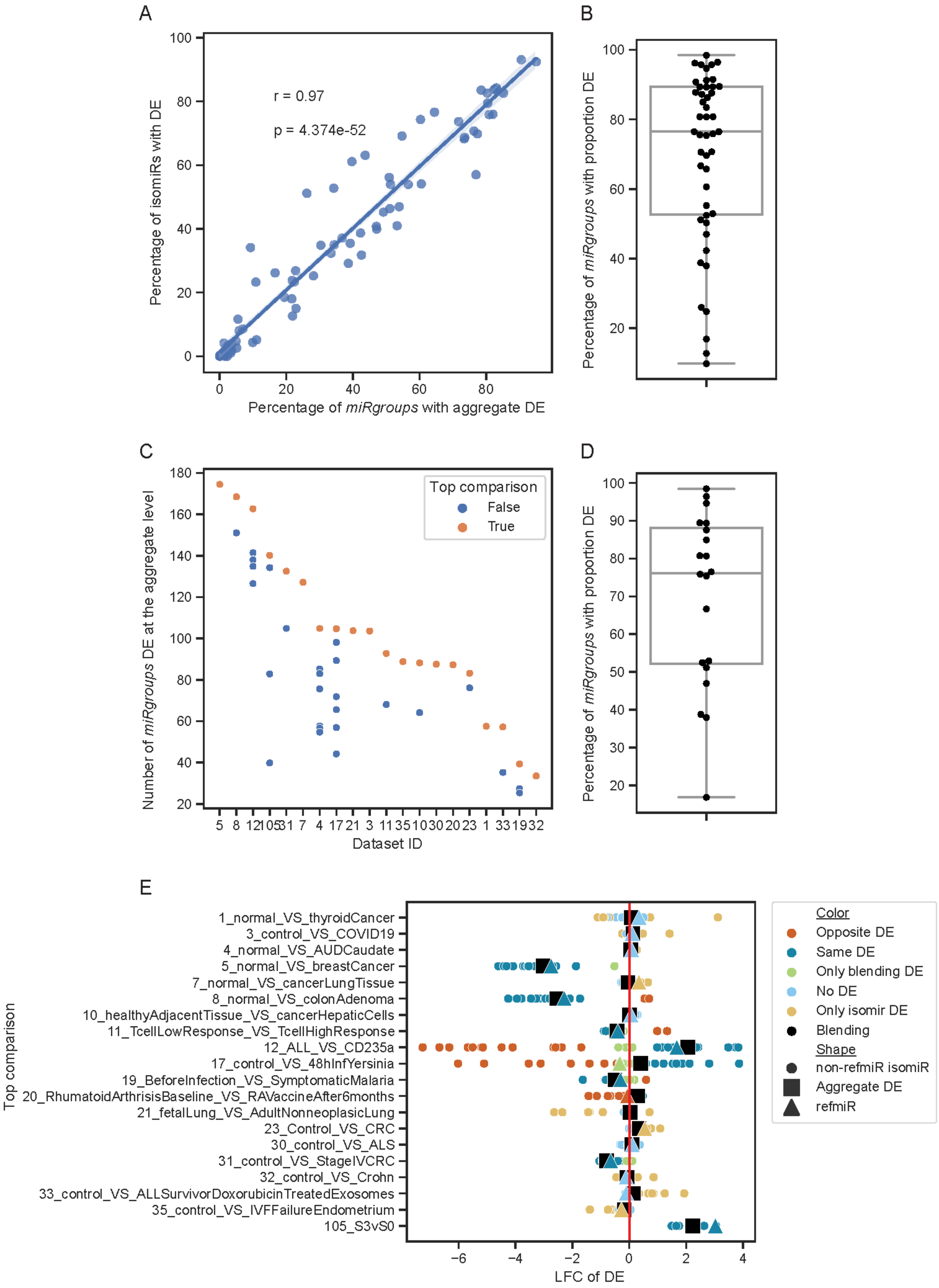

A
r = 0.97
p = 4.374e-52
Percentage of isomiRs with DE
Percentage of *miRgroups* with aggregate DE
B
Percentage of *miRgroups* with proportion DE
C
Top comparison
False
True
Number of *miRgroups* DE at the aggregate level
Dataset ID
D
Percentage of *miRgroups* with proportion DE
E
1_normal_VS_thyroidCancer
3_control_VS_COVID19
4_normal_VS_AUDCaudate
5_normal_VS_breastCancer
7_normal_VS_cancerLungTissue
8_normal_VS_colonAdenoma
10_healthyAdjacentTissue_VS_cancerHepaticCells
11_TcellLowResponse_VS_TcellHighResponse
12_ALL_VS_CD235a
17_control_VS_48hInfYersinia
19_BeforeInfection_VS_SymptomaticMalaria
20_RhumatoidArthrisisBaseline_VS_RAVaccineAfter6months
21_fetalLung_VS_AdultNonneoplasicLung
23_Control_VS_CRC
30_control_VS_ALS
31_control_VS_StageIVCRC
32_control_VS_Crohn
33_control_VS_ALLSurvivorDoxorubicinTreatedExosomes
35_control_VS_IVFFailureEndometrium
105_S3vS0
Top comparison
LFC of DE
Color
Opposite DE
Same DE
Only blending DE
No DE
Only isomir DE
Blending
Shape
non-refmiR isomiR
Aggregate DE
refmiR

**Figure S2**. **Differential expression analysis of isomiR levels**. **A**. Correlation plot showing the percentage of significantly DE isomiRs versus *miRgroups* in case-control comparisons. **B**. Percentage distribution of *miRgroups* with significant distribution changes in each case-control comparison. A *miRgroup* is flagged for change if any of its isomiRs does. **C**. Highlighting the top case-control comparison for each dataset based on the number of significantly DE *miRgroups*. These top comparisons are indicated in orange. **D**. For each selected top case-control comparison per dataset, distribution of the percentage of *miRgroups* with significant distribution shifts. **E**. DEA for miR-92-3p across top case-control comparisons in every dataset. Results for *aggregate quantification* (black square) and *isomiR quantification* (dots) are displayed. The *refmiR* is shown as a triangle. Dot and triangle colorations represent *aggregate* vs *isomiR* DE comparison classification. The x-axis indicates the LFC for each comparison.

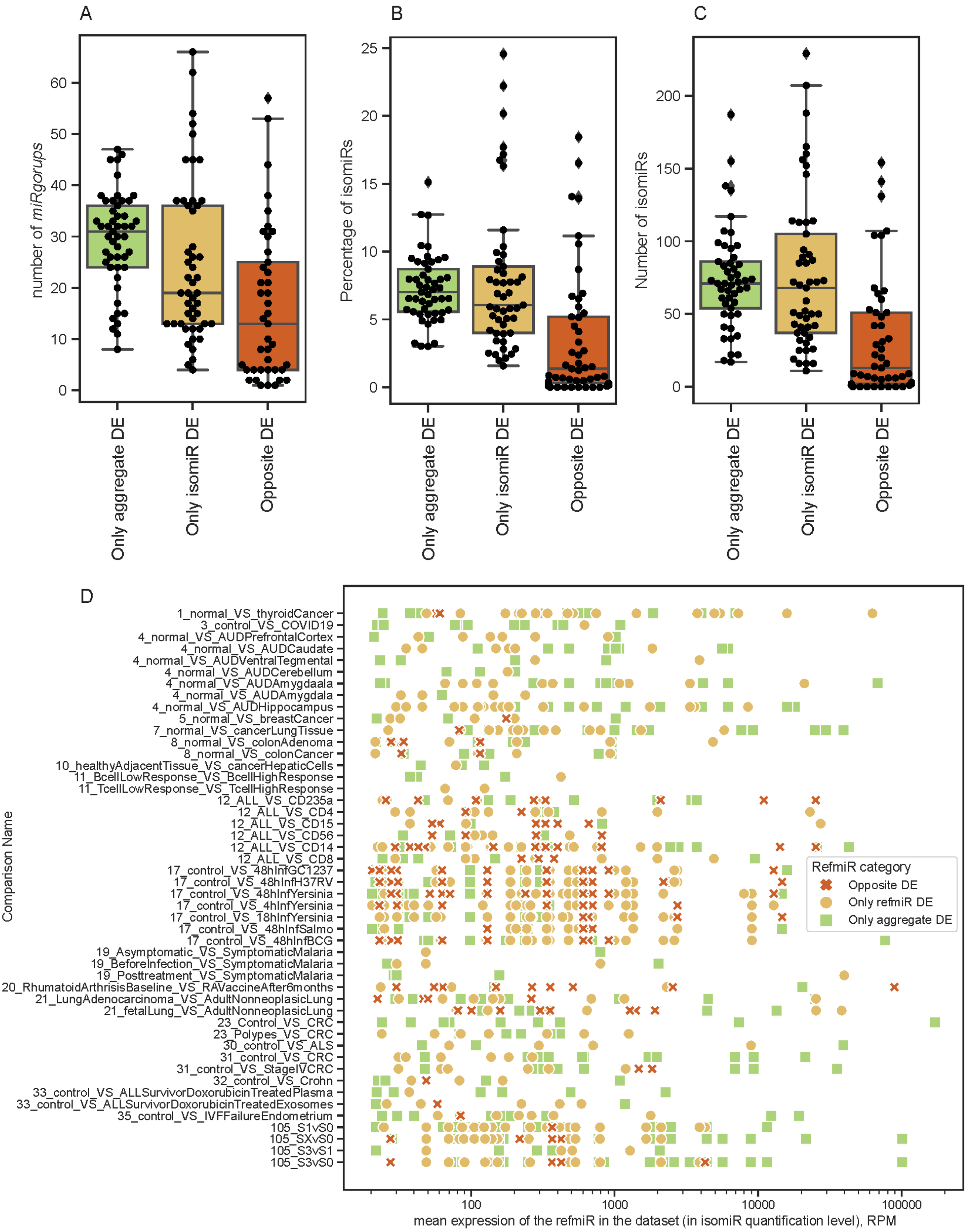
A
number of *miRgorups*
0
10
20
30
40
50
60
Only aggregate DE
Only isomiR DE
Opposite DE
B
Percentage of isomiRs
0
5
10
15
20
25
Only aggregate DE
Only isomiR DE
Opposite DE
C
Number of isomiRs
0
50
100
150
200
Only aggregate DE
Only isomiR DE
Opposite DE
D
Comparison Name
1_normal_VS_thyroidCancer
3_control_VS_COVID19
4_normal_VS_AUDPrefrontalCortex
4_normal_VS_AUDCaudate
4_normal_VS_AUDVentralTegmental
4_normal_VS_AUDCerebellum
4_normal_VS_AUDAmygdaala
4_normal_VS_AUDAmygdala
4_normal_VS_AUDHippocampus
5_normal_VS_breastCancer
7_normal_VS_cancerLungTissue
8_normal_VS_colonAdenoma
8_normal_VS_colonCancer
10_healthyAdjacentTissue_VS_cancerHepaticCells
11_BcellLowResponse_VS_BcellHighResponse
11_TcellLowResponse_VS_TcellHighResponse
12_ALL_VS_CD235a
12_ALL_VS_CD4
12_ALL_VS_CD15
12_ALL_VS_CD56
12_ALL_VS_CD14
12_ALL_VS_CD8
17_control_VS_48hInfGC1237
17_control_VS_48hInfH37RV
17_control_VS_48hInfYersinia
17_control_VS_4hInfYersinia
17_control_VS_18hInfYersinia
17_control_VS_48hInfSalmo
17_control_VS_48hInfBCG
19_Asymptomatic_VS_SymptomaticMalaria
19_BeforeInfection_VS_SymptomaticMalaria
19_Posttreatment_VS_SymptomaticMalaria
20_RhumatoidArthrisisBaseline_VS_RAVaccineAfter6months
21_LungAdenocarcinoma_VS_AdultNonneoplasicLung
21_fetalLung_VS_AdultNonneoplasicLung
23_Control_VS_CRC
23_Polypes_VS_CRC
30_control_VS_ALS
31_control_VS_CRC
31_control_VS_StageIVCRC
32_control_VS_Crohn
33_control_VS_ALLSurvivorDoxorubicinTreatedPlasma
33_control_VS_ALLSurvivorDoxorubicinTreatedExosomes
35_control_VS_IVFFailureEndometrium
105_S1vS0
105_SXvS0
105_S3vS1
105_S3vS0
100
1000
10000
100000
mean expression of the refmiR in the dataset (in isomiR quantification level), RPM
RefmiR category
Opposite DE
Only refmiR DE
Only aggregate DE

**Figure S3**. **Detailed distribution of discrepancies in differential expression patterns, between *isomiR* and *aggregate quantification*. A-C.** Distribution of the three discrepancy categories, *Only aggregate DE*, *Only isomiR DE* and *Opposite DE*, in terms of number of *miRgroups* (**A**), percentage of isomiRs (**B**) and total number of isomiRs (**C**) across all case-control comparisons. **D**. Distribution of *refmiR* DE discrepancies between *aggregate* and *isomiR quantification*. Each shape represents a specific *refmiR* with a discrepancy, and the nature of the discrepancy is denoted by both its shape and color. The x-axis reflects the mean expression of the *refmiR* in the dataset (in RPM), and the y-axis identifies the particular case-control comparison being analyzed.

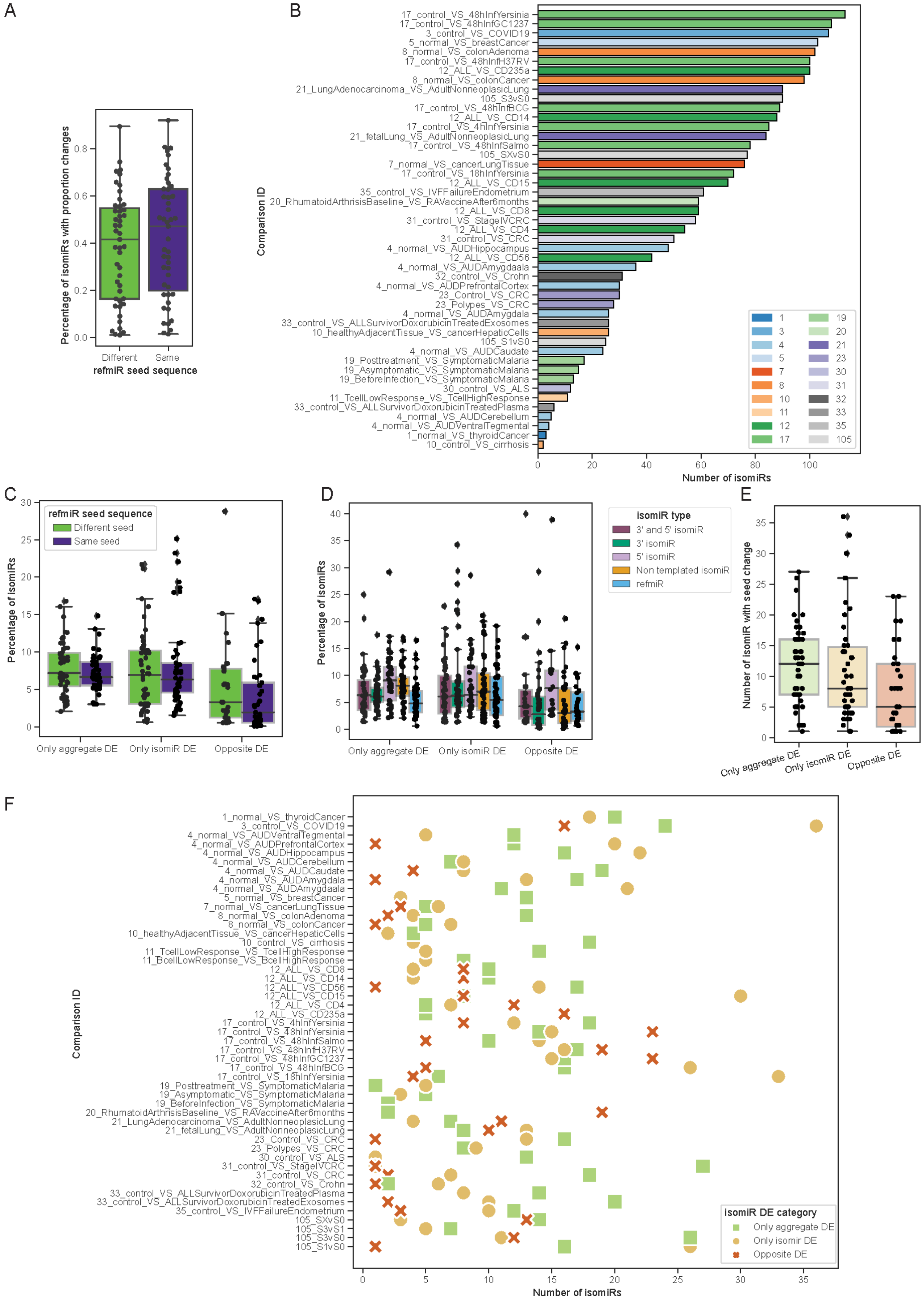
A
Percentage of isomiRs with proportion changes
0.0
0.2
0.4
0.6
0.8
Different
Same
refmiR seed sequence
B
Comparison ID
17_control_VS_48hInfYersinia
17_control_VS_48hInfGC1237
3_control_VS_COVID19
5_normal_VS_breastCancer
8_normal_VS_colonAdenoma
17_control_VS_48hInfH37RV
12_ALL_VS_CD235a
8_normal_VS_colonCancer
21_LungAdenocarcinoma_VS_AdultNonneoplasicLung
105_S3vS0
17_control_VS_48hInfBCG
12_ALL_VS_CD14
17_control_VS_4hInfYersinia
21_fetalLung_VS_AdultNonneoplasicLung
17_control_VS_48hInfSalmo
105_SXvS0
7_normal_VS_cancerLungTissue
17_control_VS_18hInfYersinia
12_ALL_VS_CD15
35_control_VS_IVFFailureEndometrium
20_RhumatoidArthrisisBaseline_VS_RAVaccineAfter6months
12_ALL_VS_CD8
31_control_VS_StageIVCRC
12_ALL_VS_CD4
31_control_VS_CRC
4_normal_VS_AUDHippocampus
12_ALL_VS_CD56
4_normal_VS_AUDAmygdaala
32_control_VS_Crohn
4_normal_VS_AUDPrefrontalCortex
23_Control_VS_CRC
23_Polypes_VS_CRC
4_normal_VS_AUDAmygdala
33_control_VS_ALLSurvivorDoxorubicinTreatedExosomes
10_healthyAdjacentTissue_VS_cancerHepaticCells
105_S1vS0
4_normal_VS_AUDCaudate
19_Posttreatment_VS_SymptomaticMalaria
19_Asymptomatic_VS_SymptomaticMalaria
19_BeforeInfection_VS_SymptomaticMalaria
30_control_VS_ALS
11_TcellLowResponse_VS_TcellHighResponse
33_control_VS_ALLSurvivorDoxorubicinTreatedPlasma
4_normal_VS_AUDCerebellum
4_normal_VS_AUDVentralTegmental
1_normal_VS_thyroidCancer
10_control_VS_cirrhosis
0
20
40
60
80
100
Number of isomiRs
1
3
4
5
7
8
10
11
12
17
19
20
21
23
30
31
32
33
35
105
C
Percentage of isomiRs
0
5
10
15
20
25
30
refmiR seed sequence
Different seed
Same seed
Only aggregate DE
Only isomiR DE
Opposite DE
D
Percentage of isomiRs
0
5
10
15
20
25
30
35
40
isomiR type
3' and 5' isomiR
3' isomiR
5' isomiR
Non templated isomiR
refmiR
Only aggregate DE
Only isomiR DE
Opposite DE
E
Number of isomiR with seed change
0
5
10
15
20
25
30
35
Only aggregate DE
Only isomiR DE
Opposite DE
F
Comparison ID
1_normal_VS_thyroidCancer
3_control_VS_COVID19
4_normal_VS_AUDVentralTegmental
4_normal_VS_AUDPrefrontalCortex
4_normal_VS_AUDHippocampus
4_normal_VS_AUDCerebellum
4_normal_VS_AUDCaudate
4_normal_VS_AUDAmygdala
4_normal_VS_AUDAmygdaala
5_normal_VS_breastCancer
7_normal_VS_cancerLungTissue
8_normal_VS_colonAdenoma
8_normal_VS_colonCancer
10_healthyAdjacentTissue_VS_cancerHepaticCells
10_control_VS_cirrhosis
11_TcellLowResponse_VS_TcellHighResponse
11_BcellLowResponse_VS_BcellHighResponse
12_ALL_VS_CD8
12_ALL_VS_CD14
12_ALL_VS_CD56
12_ALL_VS_CD15
12_ALL_VS_CD4
12_ALL_VS_CD235a
17_control_VS_4hInfYersinia
17_control_VS_48hInfYersinia
17_control_VS_48hInfSalmo
17_control_VS_48hInfH37RV
17_control_VS_48hInfGC1237
17_control_VS_48hInfBCG
17_control_VS_18hInfYersinia
19_Posttreatment_VS_SymptomaticMalaria
19_Asymptomatic_VS_SymptomaticMalaria
19_BeforeInfection_VS_SymptomaticMalaria
20_RhumatoidArthrisisBaseline_VS_RAVaccineAfter6months
21_LungAdenocarcinoma_VS_AdultNonneoplasicLung
21_fetalLung_VS_AdultNonneoplasicLung
23_Control_VS_CRC
23_Polypes_VS_CRC
30_control_VS_ALS
31_control_VS_StageIVCRC
31_control_VS_CRC
32_control_VS_Crohn
33_control_VS_ALLSurvivorDoxorubicinTreatedPlasma
33_control_VS_ALLSurvivorDoxorubicinTreatedExosomes
35_control_VS_IVFFailureEndometrium
105_SXvS0
105_S3vS1
105_S3vS0
105_S1vS0
0
5
10
15
20
25
30
35
Number of isomiRs
isomiR DE category
Only aggregate DE
Only isomir DE
Opposite DE

**Figure S4. DE patterns of isomiRs with seed sequence change. A**. Distribution of the percentage of isomiRs with significant proportion changes, comparing those with and without a seed sequence change. **B**. Count of isomiRs displaying both seed change and significant differential distribution across case-control comparisons. Bars are color-coded by dataset ID. **C**. Distribution of isomiRs with discrepancies between *aggregate* and *isomiR quantification*, stratified by seed sequence change, across case-control comparisons. **D**. Distribution of isomiRs with discrepancies between *aggregate* and *isomiR quantification*, categorized by isomiR type, across case-control comparisons. **E**. Tally of isomiRs with a seed change and discrepancies between *aggregate* and *isomiR quantification*, classified by discrepancy type, across all case-control comparisons. **F**. Analysis of isomiRs with seed changes displaying DE discrepancies. The y-axis represents the case-control comparison, the x-axis indicates the count of isomiRs with discrepancies, and shapes and colors differentiate discrepancy categories.

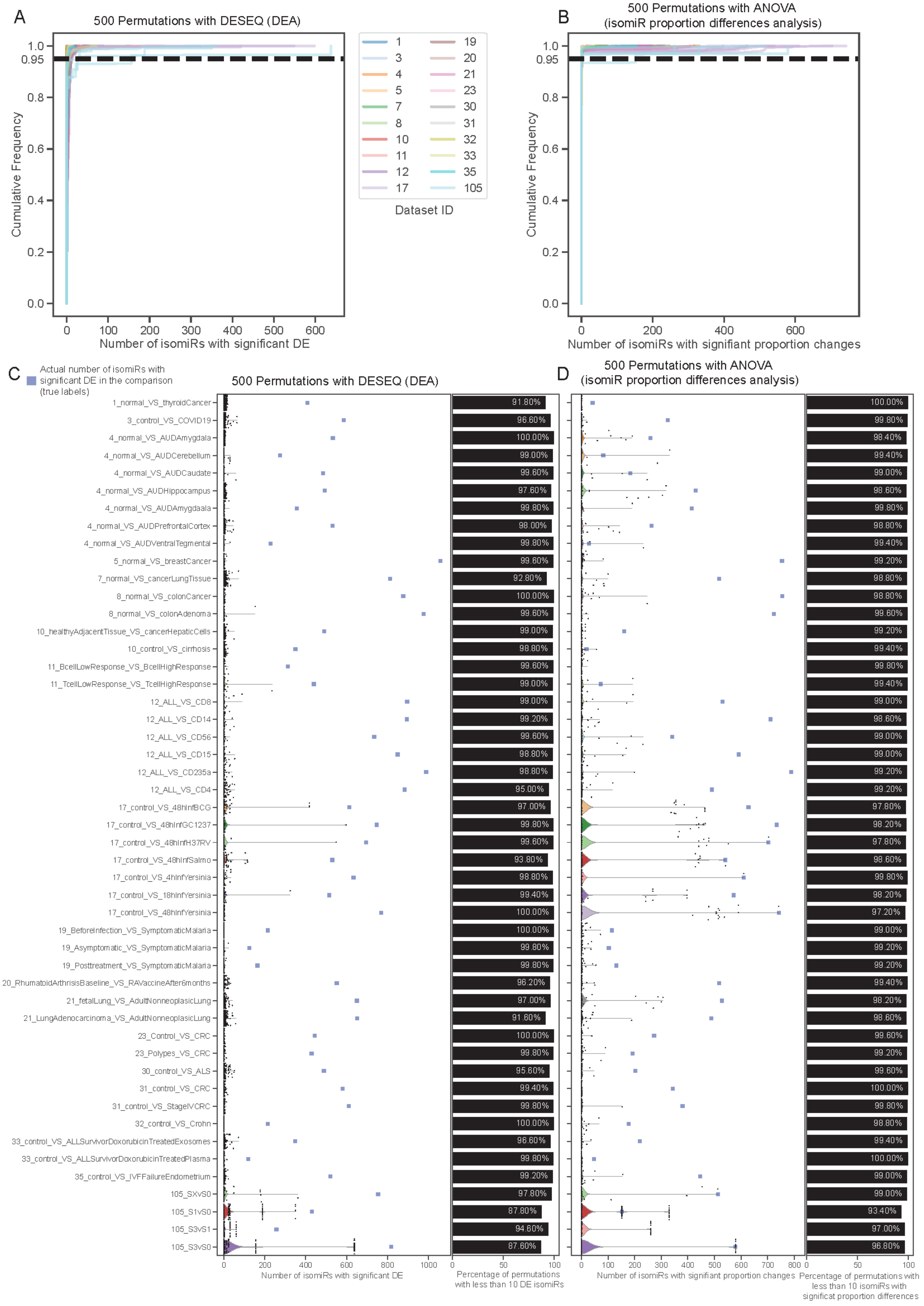

A
500 Permutations with DESEQ (DEA)
Cumulative Frequency
Number of isomiRs with significant DE
Dataset ID
1
3
4
5
7
8
10
11
12
17
19
20
21
23
30
31
32
33
35
105
B
500 Permutations with ANOVA
(isomiR proportion differences analysis)
Cumulative Frequency
Number of isomiRs with signifiant proportion changes
C
Actual number of isomiRs with significant DE in the comparison (true labels)
500 Permutations with DESEQ (DEA)
D
500 Permutations with ANOVA
(isomiR proportion differences analysis)
1_normal_VS_thyroidCancer
3_control_VS_COVID19
4_normal_VS_AUDAmygdala
4_normal_VS_AUDCerebellum
4_normal_VS_AUDCaudate
4_normal_VS_AUDHippocampus
4_normal_VS_AUDAmygdaala
4_normal_VS_AUDPrefrontalCortex
4_normal_VS_AUDVentralTegmental
5_normal_VS_breastCancer
7_normal_VS_cancerLungTissue
8_normal_VS_colonCancer
8_normal_VS_colonAdenoma
10_healthyAdjacentTissue_VS_cancerHepaticCells
10_control_VS_cirrhosis
11_BcellLowResponse_VS_BcellHighResponse
11_TcellLowResponse_VS_TcellHighResponse
12_ALL_VS_CD8
12_ALL_VS_CD14
12_ALL_VS_CD56
12_ALL_VS_CD15
12_ALL_VS_CD235a
12_ALL_VS_CD4
17_control_VS_48hInfBCG
17_control_VS_48hInfGC1237
17_control_VS_48hInfH37RV
17_control_VS_48hInfSalmo
17_control_VS_4hInfYersinia
17_control_VS_18hInfYersinia
17_control_VS_48hInfYersinia
19_BeforeInfection_VS_SymptomaticMalaria
19_Asymptomatic_VS_SymptomaticMalaria
19_Posttreatment_VS_SymptomaticMalaria
20_RhumatoidArthrisisBaseline_VS_RAVaccineAfter6months
21_fetalLung_VS_AdultNonneoplasicLung
21_LungAdenocarcinoma_VS_AdultNonneoplasicLung
23_Control_VS_CRC
23_Polypes_VS_CRC
30_control_VS_ALS
31_control_VS_CRC
31_control_VS_StageIVCRC
32_control_VS_Crohn
33_control_VS_ALLSurvivorDoxorubicinTreatedExosomes
33_control_VS_ALLSurvivorDoxorubicinTreatedPlasma
35_control_VS_IVFFailureEndometrium
105_SXvS0
105_S1vS0
105_S3vS1
105_S3vS0
91.80%
96.60%
100.00%
99.00%
99.60%
97.60%
99.80%
98.00%
99.80%
99.60%
92.80%
100.00%
99.60%
99.00%
98.80%
99.60%
99.00%
99.00%
99.20%
99.60%
98.80%
98.80%
95.00%
97.00%
99.80%
99.60%
93.80%
98.80%
99.40%
100.00%
100.00%
99.80%
99.80%
96.20%
97.00%
91.60%
100.00%
99.80%
95.60%
99.40%
99.80%
100.00%
96.60%
99.80%
99.20%
97.80%
87.80%
94.60%
87.60%
Number of isomiRs with significant DE
Percentage of permutations with less than 10 DE isomiRs
100.00%
99.80%
98.40%
99.40%
99.00%
98.60%
99.80%
98.80%
99.40%
99.20%
98.80%
98.80%
99.60%
99.20%
99.40%
99.80%
99.40%
99.00%
98.60%
99.00%
99.00%
99.20%
99.20%
97.80%
98.20%
97.80%
98.60%
99.80%
98.20%
97.20%
99.00%
99.20%
99.20%
99.40%
98.20%
98.60%
99.60%
99.20%
99.60%
100.00%
99.80%
98.80%
99.40%
100.00%
99.00%
99.00%
93.40%
97.00%
96.80%
Number of isomiRs with signifiant proportion changes
Percentage of permutations with less than 10 isomiRs with significat proportion differences

**Figure S5. Permutation analysis for proportion changes and DEA. A-B.** Cumulative frequency curve for the 500 permutation of sample labels for DEA (**A**) and proportion changes (**B**). For each comparison, the plotted line represents the cumulative proportion of permutation with the number of isomiRs with statistically significant changes represented on the x axis. The lines are colored by dataset. The dashed line represents 95% of permutations (for alpha = 0.05). **C-D.** Breakdown of the permutation analysis results, by comparison, for DEA (**C**) and for proportion changes (**D**). For each comparison a violin plot and a stripplot represent the 500 permutation results (black dots). The x axis represents the number of isomiRs with statistically significant changes. The blue squares represent the actual results with the true labels. On the right, a bar plot shows the percentage of permutations with less than 10 isomiRs with significant changes.

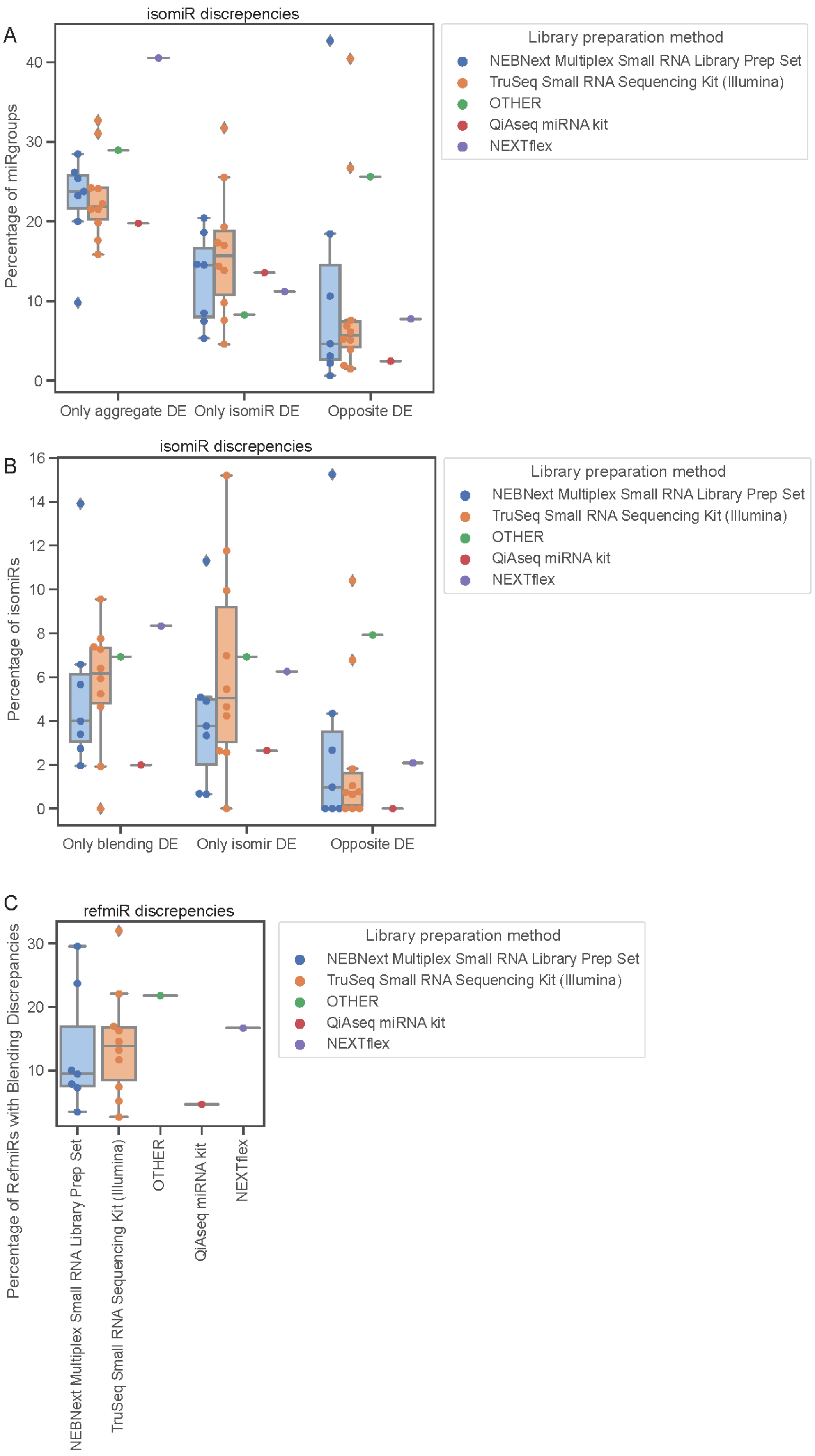
A
isomiR discrepencies
Percentage of miRgroups
0
10
20
30
40
Only aggregate DE
Only isomiR DE
Opposite DE
Library preparation method
NEBNext Multiplex Small RNA Library Prep Set
TruSeq Small RNA Sequencing Kit (Illumina)
OTHER
QiAseq miRNA kit
NEXTflex
B
isomiR discrepencies
Percentage of isomiRs
0
2
4
6
8
10
12
14
16
Only blending DE
Only isomir DE
Opposite DE
Library preparation method
NEBNext Multiplex Small RNA Library Prep Set
TruSeq Small RNA Sequencing Kit (Illumina)
OTHER
QiAseq miRNA kit
NEXTflex
C
refmiR discrepencies
Percentage of RefmiRs with Blending Discrepancies
10
20
30
NEBNext Multiplex Small RNA Library Prep Set
TruSeq Small RNA Sequencing Kit (Illumina)
OTHER
QiAseq miRNA kit
NEXTflex
Library preparation method
NEBNext Multiplex Small RNA Library Prep Set
TruSeq Small RNA Sequencing Kit (Illumina)
OTHER
QiAseq miRNA kit
NEXTflex

**Figure S6. Impact of library preparation methods in quantification discrepancies. A-B.** Percentage of *miRgroups* (**A**) and isomiRs (**B**) that show discrepancies between *aggregate* and *isomiR quantification*, separated by library preparation method. Here only the top comparison for each dataset is selected. **C.** Representation of all discrepancies between refmiR and aggregate signals, presented as a percentage of refmiRs, separated by library preparation method.

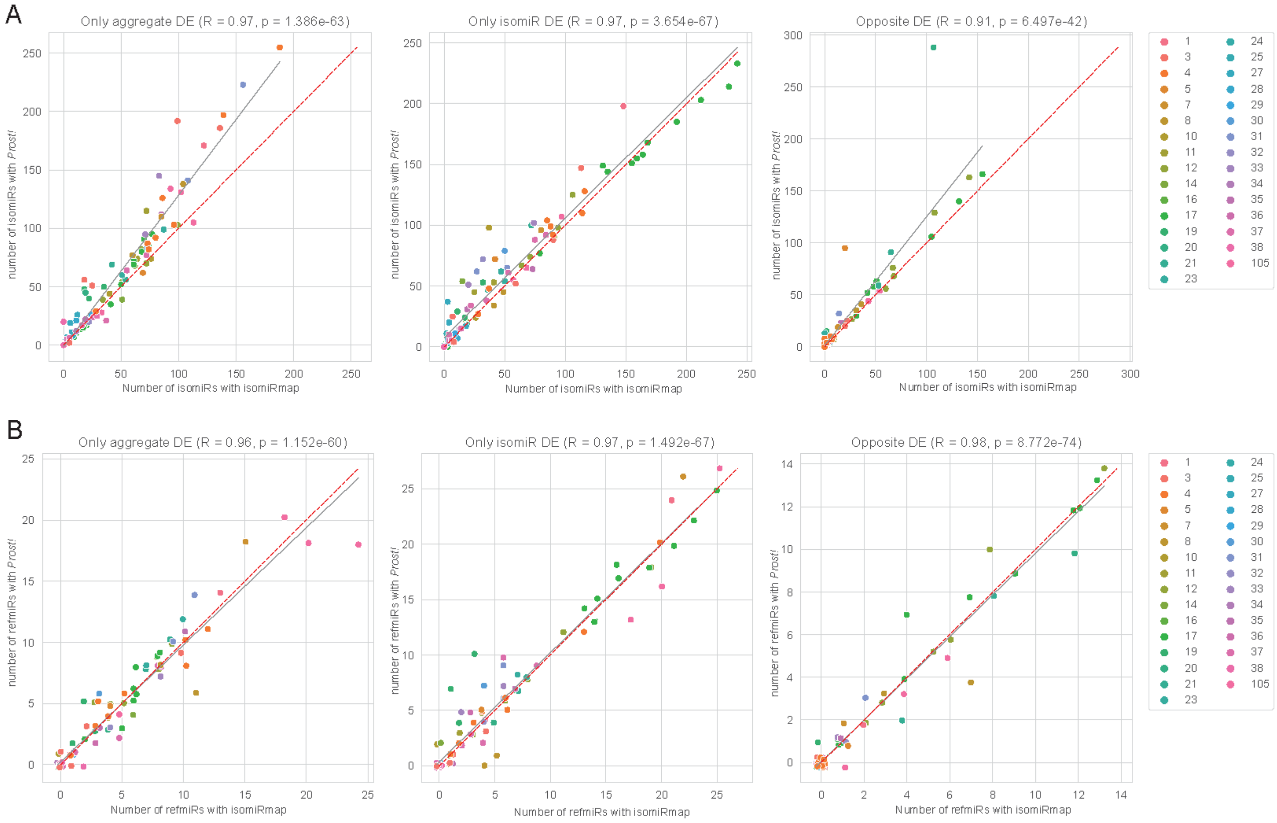


**Figure S7. Comparison of isomiRmap and *Prost!* isomiR analysis results.** Cases of discrepancies were counted between *aggregate* and *isomiR quantification*, when *isomiR quantification* was performed by isomiRmap (x axis) and *Prost!* (y axis). The dashed red line represents a 1/1 curve, and a correlation fitted curve by a gray line. Panel **A** represents all isomiRs, and Panel **B** concentrates on the discrepancies in *refmiRs*.

### Supplementary Tables

**Supplementary Table 1: List and references of the 28 publicly available datasets used in this study.** All datasets are available on GEO, and were downloaded through SRA from their GEO page. Sample - level information was obtained from the SRA annotation tables.

**Supplementary Table 2: Detail on the Case - Control comparisons run on the 28 publicly available datasets and the generated hypoxia compartment dataset.** Columns indicated the dataset id (dataset 104 corresponds to the new hypoxia-compartment dataset) and the comparison name. Samples are selected to belong in the case or control group for DEA based on their annotations in the SRA sample annotation tables. The corresponding column names and values associated are described.
